# Cerebral Dopamine Neurotrophic Factor regulates multiple neuronal subtypes and behavior

**DOI:** 10.1101/733949

**Authors:** Yu-Chia Chen, Diego Baronio, Svetlana Semenova, Shamsiiat Abdurakhmanova, Pertti Panula

**Affiliations:** Department of Anatomy, University of Helsinki, Finland

## Abstract

Cerebral Dopamine Neurotrophic Factor (CDNF) protects dopaminergic neurons against toxic damage in the rodent brain, and is in clinical trials to treat Parkinson’s disease patients. Yet the underlying mechanism is poorly understood. To examine its mode of action and significance, we examined the development of neurotransmitter systems from larval to adult mutant zebrafish lacking *cdnf*. Although a lack of *cdnf* did not affect overall brain dopamine levels, dopaminergic neuronal clusters showed significant abnormalities. The number of histamine neurons that surround the dopaminergic neurons was significantly reduced. Expression of *tyrosine hydroxylase 2* in the brain was elevated in *cdnf* mutants throughout their lifespan. There were abnormally few GABA neurons in the hypothalamus in the mutant larvae, and expression of glutamate decarboxylase was reduced throughout the brain. *cdnf* mutant adults showed a range of behavioral phenotypes, including increased sensitivity to pentylenetetrazole-induced seizures. Shoaling behavior of mutant adults was abnormal, and they did not display social attraction to conspecifics. CDNF plays a profound role in shaping the neurotransmitter circuit structure, seizure susceptibility, and complex behaviors in zebrafish. These findings are informative for dissecting the diverse functions of this poorly understood factor in human conditions related to Parkinson’s disease and complex behaviors

## Introduction

Neurotrophic factors (NTFs), such as neurotrophins, glial cell line-derived neurotrophic factor family of ligands, and neurokines are crucial regulators of neurogenesis and regeneration. These secretory proteins and their signaling receptors are responsible for the survival, maintenance, and synaptic plasticity of nervous systems through development to adulthood (1). Several neurodegenerative disorders such as Parkinson’s disease (PD) and Alzheimer’s disease (AD) are associated with dysregulation of trophic factors (2, 3).

An unconventional NTF family, which has a distinct two-domain protein structure and trophic effects on dopaminergic neurons, has recently been identified (4, 5). This evolutionarily conserved NTF family contains two vertebrate proteins – mesencephalic astrocyte-derived neurotrophic factor (MANF) and cerebral dopamine neurotrophic factor (CDNF) (6). CDNF and MANF both protect dopaminergic neurons against oxidative stress, neurotoxins, cerebral ischemia, and neuroinflammation-induced neuronal death (6–8). As such, CDNF has become recognized as a promising candidate for clinical treatment of PD due to its potent neuroprotective and neurorestorative effects on midbrain dopamine neurons (6, 9, 10).

CDNF and MANF are widely distributed in the mammalian brain and peripheral organs (11, 12). Their protein structure contains two main functional motifs. One is the N-terminus, which is similar to the saposin-like domain that has the capacity of lipid/cell membrane binding. The other is the C-terminus, composed of the unfolded Cys-X-X-Cys (CXXC) motif, the SAP domain of Ku70, and a putative endoplasmic reticulum (ER) retention signal (KDEL/RTDL) at the end of the C-terminal, which may protect cells from ER stress-induced apoptosis (4, 5, 13). In addition to these two functional domains, the eight helices structure may function in protein-protein interactions.

For example, in *Drosophila melanogaster* and *Caenorhabditis elegans*, loss of MANF causes embryonic lethality and deficiencies in dopaminergic neurons (14, 15). In zebrafish, *manf* depletion also causes a reduction of dopaminergic neurons in the brain (16). Although such a dopaminergic phenotype in the CNS is not observed in MANF knock-out mice, they do have a diabetic syndrome, possibly due to increased ER stress caused by reduced beta cell proliferation (17, 18).

CDNF protects cultured mesencephalic neurons against alpha-synuclein-oligomer-induced toxicity (5). In 6-hydroxydopamine (6-OHDA) and 1-methyl-4-phenyl-1,2,3,6-tetrahydropyridine (MPTP)-induced Parkinsonian animal models, the application of CDNF and MANF protects and rescues midbrain dopamine neurons (11, 19–21). In addition to its neuroprotective effects in PD animal models, CDNF has been shown to improve long-term memory in an APP/PS1 mouse model of AD (22), and reduce Aß25-35-induced ER stress and synaptotoxicity in cultured hippocampus neurons (23). In non-neuronal cells, CDNF improves cell viability and protects cultured cardiomyocytes from apoptosis induced by ER stress (24). Growing evidence suggests that MANF and CDNF are ER stress response proteins involved in the unfolded protein response (UPR) through interactions with glucose-regulated protein 78 (BiP/GRP78) (25, 26). Nevertheless, how CDNF responds to ER stress and its functions under healthy conditions remain largely unknown. The lack of effect of CDNF on normal cells is in striking contrast with its effect on the lesioned brain (6).

To date, the role of CDNF in the development of dopaminergic or other neurons has not been addressed. As such, its role and mechanisms of action have remained elusive. To examine the biological function of cdnf, we first generated zebrafish null mutants by CRISPR/Cas9 genome editing. We investigated the role of cdnf on major neurotransmitter systems in the CNS, including dopaminergic, histaminergic, serotonergic, and GABAergic circuits using qPCR, *in situ* hybridization, immunohistochemistry, and HPLC analysis, as well as locomotor behavioral analysis, from development throughout their life span. Remarkably, we found that adult cdnf mutant zebrafish show abnormal social behaviors and seizure susceptibility phenotypes that are conceivably associated with the multiple impairments of major neurotransmitter networks. These results link CDNF and the UPR to complex behaviors, which may have important implications for human neurological disorders.

## Results

### Zebrafish cdnf 3D structure and mRNA distribution during embryogenesis and in adult tissues

Currently there is one human homologous *cdnf* (NM_001123281.1) documented in the latest zebrafish database (GRCz11). The zebrafish *cdnf* gene is located on chromosome 4 and contains four exons with the exon-intron splice sites conserved in mammalian CDNF and MANF. The open reading frame encodes a protein of 182 amino acid residues with a 25 amino-acid signal peptide, and shares 67.4% and 65.1% of amino acid sequence similarity with zebrafish manf (NP_001070097) and human CDNF (NP_001025125), respectively. The predicted secondary structure analyzed by the protein homology/analogy recognition engine Phyre2 (27) indicated that zebrafish cdnf contained seven α-helices (Figure 1A) and eight positioned-conserved cysteine residues. The N-terminus domain is highly similar to human saposin D, and the C-terminus is analogous to the SAP domain (Figure 1B), suggesting that zebrafish cdnf is structurally conserved with human CDNF (13). To determine the spatiotemporal expression of *cdnf*, WISH and qPCR was performed in embryos and adult tissues. The *cdnf* mRNA was widely expressed at 1 dpf in the brain, eyes, and muscles. In 2- and 3-dpf larvae, *cdnf* transcripts were detected in the midbrain-hindbrain boundary, hindbrain, otic vesicles, and heart (Figure 1C). qPCR detected *cdnf* transcripts as early as 2 hpf (Figure 1D), and expression gradually increased after 1 dpf. In adult organs, the highest expression level was found in the eyes compared with the brain, kidney, and liver; no sex differences were detected (Figure 1E).

**Figure 1.**
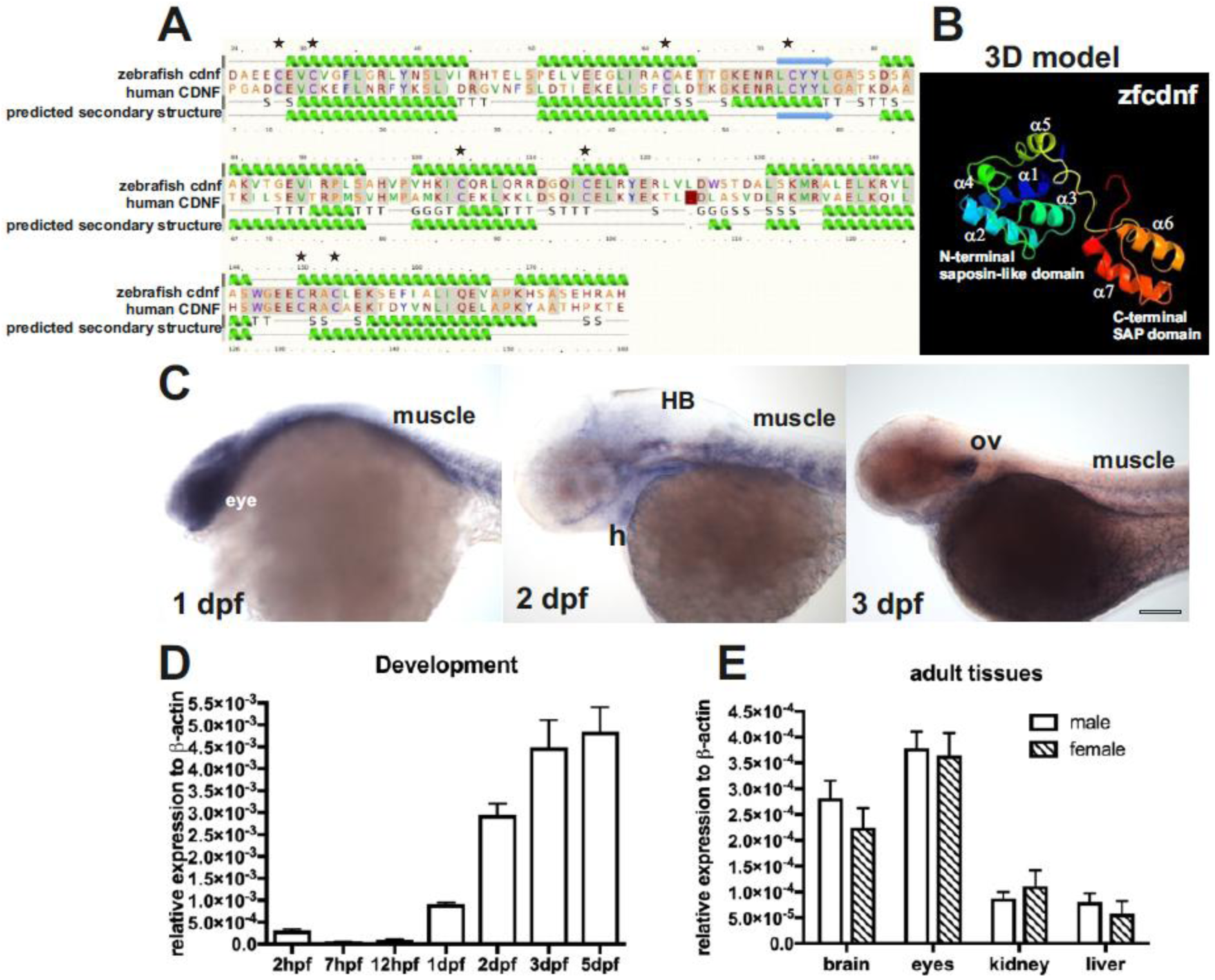
Predicted domain structures and spatiotemporal expression of zebrafish cdnf. (**A**) Secondary structure, and (**B**) predictive 3D model of zebrafish cdnf by Phyre2 modeling with human CDNF. (**C**) whole-mount *in situ* hybridization highlighting *cdnf* expression in the head, eyes, heart, muscles, and optic vesicles during development. (**D**) quantitative RT-PCR showing the zygotic *cdnf* mRNA expression at various developmental stages (n=3), and (**E**) in various adult organs (n=4). Green helices indicate alpha-helices, blue arrows indicate beta-strands. Grey lines indicate the conserved amino acid residues. G indicates the 3-trun helix. T indicates hydrogen bonded turn. S indicates the bend. H, heart. HB, hindbrain. ov, optic vesicle. Data are mean ± SEM; one-way ANOVA was used for statistics.

### Characterization of the CRISPR/Cas9-generated zebrafish *cdnf* mutant allele

To study the biological function of *cdnf*, we generated mutations using the CRISPR/Cas9 system. A sequence-specific guide RNA was designed to target exon2 of *cdnf* (Figure 2A). We identified two reading frame-shift mutant alleles: one with a 14 base-pair deletion (Figure 2A and 2B) and another with a 1 base-pair insertion (data not shown). Both lesions are located in exon2, causing premature termination of the protein after amino acid 49 (NP_001116753.1, Figure 2B). We also cloned and sequenced the full-length reading frame in *cdnf* transcripts isolated from the mutant brain. The deletion sequence was the same as the genomic deletion sequence. In this study, the *cdnf* 14 base-pair deletion mutant allele was used for subsequent experiments. Heterozygous *cdnf* mutants (F3 and later generations) were mated to generate wild-type (WT), heterozygous (HET), and homozygous mutant embryos. The ratio of genotyped sibling offspring matched the normal Mendelian ratio (1:2:1). All progenies were tail clipped at 3 dpf and genotyped using HRM analysis, according to the distinct melting curves of each genotype. The embryos with the same HRM curve were grouped together before 5 dpf (Figure 2C). *In situ* hybridization results revealed that *cdnf* expression was mostly abolished in the caudal raphe, ventral part of the posterior tuberculum, and tectum opticum of *cdnf* mutant brains (Figure 2D, 2E). The faint signal in the *cdnf* mutant brain may be due to the remaining truncated mRNA. The *cdnf* mRNA quantification is shown in Figure 3A-3C. *cdnf* homozygous mutants were viable, swam normally, were fertile, and had no gross morphological phenotype (Figure 2F).

**Figure 2.**
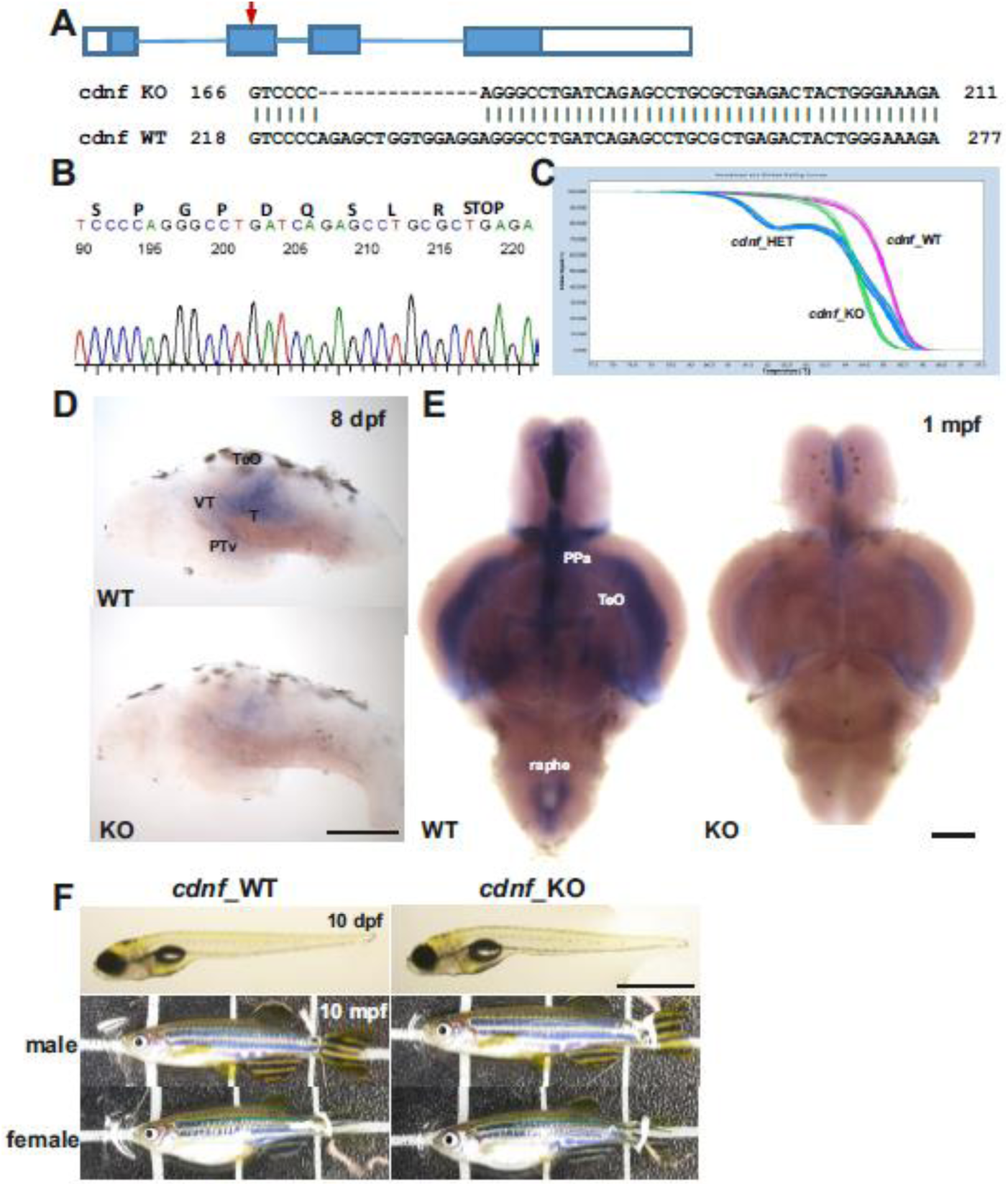
CRISPR/Cas9-generated *cdnf* mutant zebrafish. (**A**) Scheme of CRISPR/Cas9-generated 14-base pair deletion in exon2 of *cdnf*. (**B**) Sequence chromatogram showing a premature stop codon of a homozygous *cdnf* mutant zebrafish. (**C**) Results of the HRM analysis showing the distinctive melting curves of each genotype. (**D**) Reduction in *cdnf* mRNA expression shown in the brains of 8-dpf and (**E**) 1-mpf *cdnf* knock-out fish. (**F**) No obvious gross phenotype in larvae and adult *cdnf* knock-out fish. PPa, anterior parvocellular preoptic nucleus. PTv, ventral part of posterior tuberculum. T, midbrain tegmentum. TeO, tectum opticum. Scale bar is 200μm in D, 1mm (10 dpf) and 1cm (10 mpf) in F.

**Figure 3.**
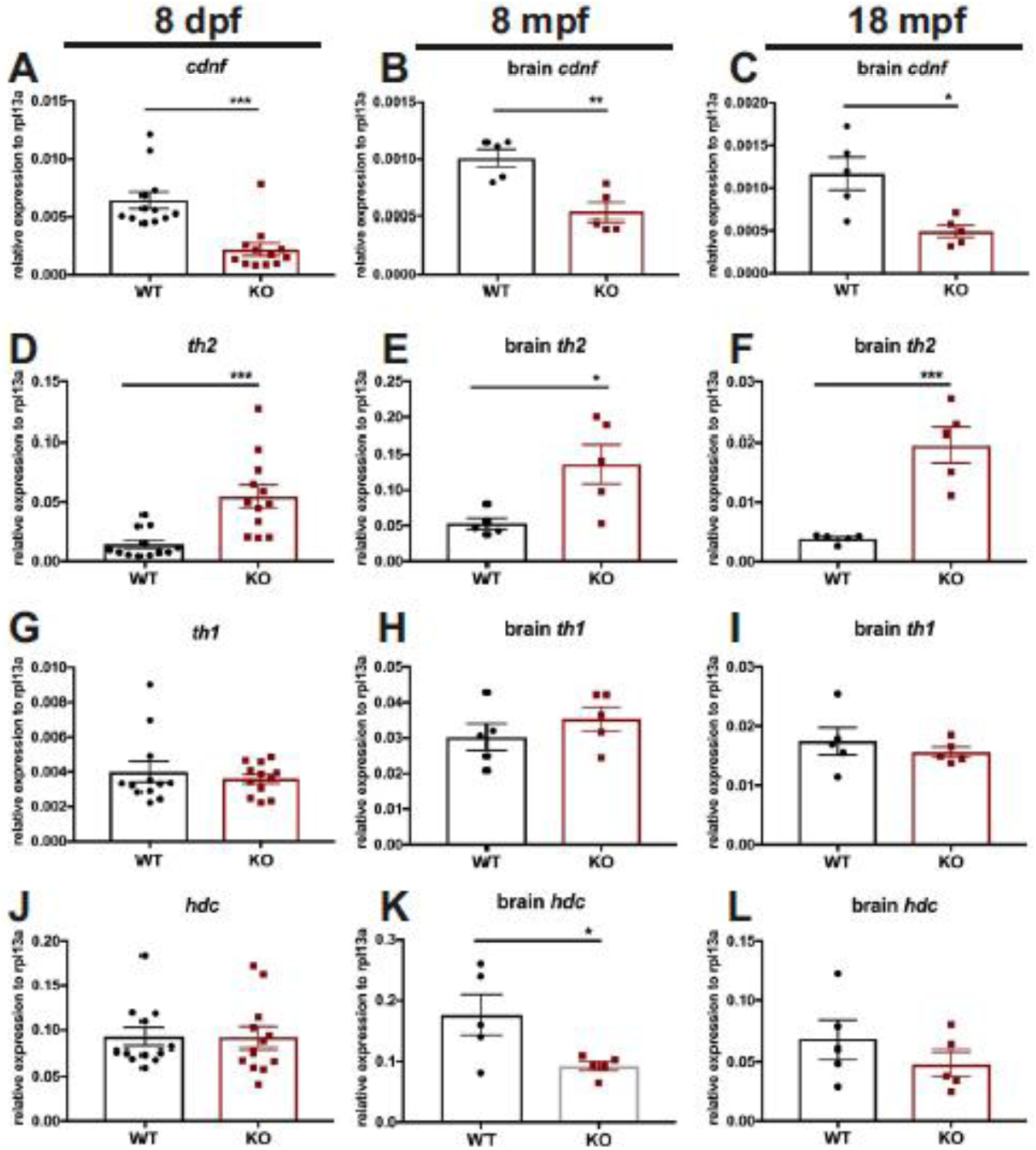
qPCR analysis of *cdnf* and neurotransmitter synthesis enzymes at larval, adulthood, and aging stages. (**A-C**) Significant reduction of *cdnf* mRNA in *cdnf*-deficient (knock-out) fish. (**D-F**) Significant increase of *th2* transcripts in *cdnf*-deficient fish. (**G-I**) Non-significant differences in *th1* mRNA expression between groups. (**J-L**), Significant reduction of *hdc* mRNA expression in adult brains. qPCR analysis, relative to expression of the housekeeping gene *rpl13a*; values are mean ± SEM. n=12 for 8-dpf fish, n=5 for 8-mpf brains, n=5 for 18-mpf brains in each *cdnf* knock-out and wild-type group. Data are mean ± SEM. Student’s *t*-test was used for statistical analysis *p<0.05, **p<0.01, and ***p<0.001.

### Upregulation of *tyrosine hydroxylase 2* (*th2*) expression in *cdnf* mutant fish

We addressed the question whether *cdnf* is required for development or maintenance of the dopaminergic neurons in the zebrafish brain. qPCR analysis was performed on 8-dpf larvae, 8-mpf brains, and 18-mpf brains to examine transcript levels of relevant marker genes of the dopaminergic and histaminergic systems (known to depend on dopaminergic system). *manf* was also analyzed to reveal if this closely related growth factor is upregulated as a consequence of genetic compensation in the *cdnf* mutant fish. In zebrafish, gene duplication has led to two non-allelic forms of human orthologous *tyrosine hydroxylase* (*th*) that are expressed in the brain in a largely complementary manner (28). We thus analyzed the mRNA levels of both tyrosine hydroxylases, *th1* and *th2*. Remarkably, a significant increase of *th2* was observed in *cdnf* mutants compared with their WT siblings (Figure 3D-3F), whereas the expression level of *th1* mRNA showed no statistically significant difference (Figure 3G-3I). Additionally, a significant downregulation of *histidine decarboxylase* (*hdc*, a histaminergic marker) transcript was observed in 8-mpf mutant brains (Figure 3J-3L). Expression of *hdc* has been shown to be regulated by dopamine produced by *th2* (29). We also confirmed that a significant reduction of *cdnf* transcript remained in *cdnf* mutants throughout their lifespan (Figure 3A-3C), which agreed with the *in situ* hybridization results (Figure 2D, 2E). Hence, 80% of the truncated *cdnf* mRNA undergoing nonsense-mediated mRNA decay pathway resulted in *cdnf*-deficient mutant fish. The mRNA level of *manf*, a closely related trophic factor, was not significantly altered (data not shown). The qPCR results revealed that *cdnf* has an impact on the regulation of dopaminergic and histaminergic gene expression.

### A dynamic change of dopaminergic neuron numbers in *cdnf* mutant fish

It is evident that dopaminergic signaling regulates the developing hypothalamic neurotransmitter identity (29). Due to the prominent upregulation of *th2* transcripts in *cdnf* mutant fish, we then quantified the number of cells of the dopaminergic populations in the prethalamus and caudal hypothalamus by immunohistochemistry to assess whether the TH1- and TH2-containing cell numbers were altered. 8-dpf dissected brains were co-stained with two antibodies that recognize dopaminergic neurons; one recognizes both TH1 and TH2 (30), and one recognizes only TH1 in zebrafish. A significant increase in TH1- and TH2-positive cell numbers was observed in TH1/TH2 group 10/10b in the caudal hypothalamus area (Hc) of knock-out (KO) mutants compared with their WT siblings (Figure 4A-4E). However, the number of TH1-positive cells was unaffected in this region (Figure 4F), suggesting that the increase in TH1- and TH2-positive cell numbers in the Hc area of *cdnf* mutant brains is due to the increase in TH2-containing neurons, consistent with the qPCR results (Figure 3D-3F) and the *in situ* hybridization result (Figure 4H). Nevertheless, a significant decrease in TH1-positive cell number was found in the prethalamus region (TH1 group 5,6,11) in *cdnf* mutants (Figure 4G). On the other hand, the serotonergic population in this location remained intact in KO mutants (Figure 4I). A dynamic change in dopaminergic neuron populations found in *cdnf* mutant fish suggests that *cdnf* may function distinctly on dopaminergic populations in different brain areas.

**Figure 4.**
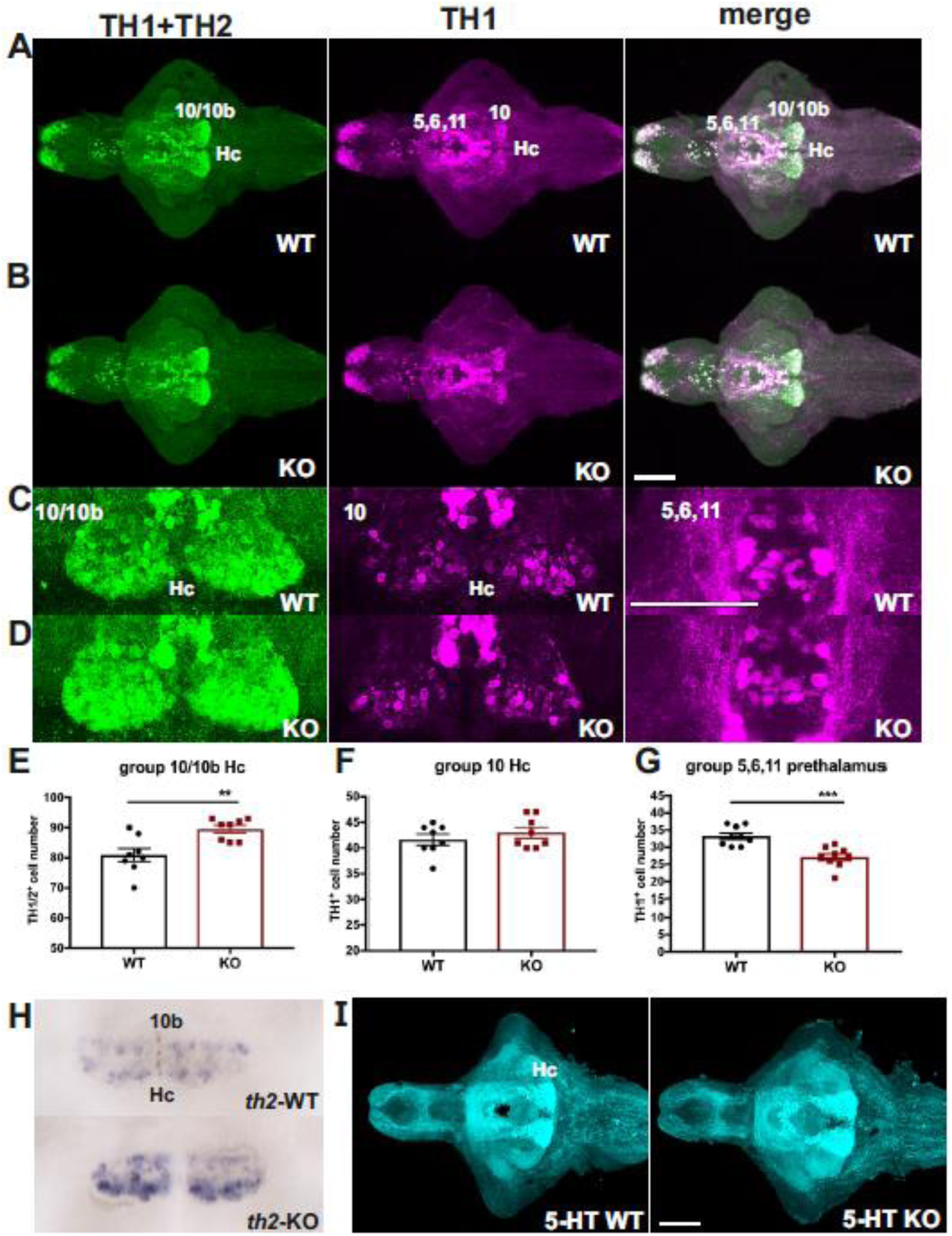
Selective alteration in dopaminergic systems in *cdnf* mutant fish. (**A**) Co-labeling of zebrafish tyrosine hydroxylase 1 (th1) and tyrosine hydroxylase 2 (th2) of 8-dpf *cdnf* wild-type brains. (**B**) Co-labeling of zebrafish tyrosine hydroxylase 1 (th1) and tyrosine hydroxylase 2 (th2) of 8-dpf *cdnf* knock-out brains. (**C**) Higher magnification images of TH2 10/10b (Hc) group, TH1 10 group, and TH1 5,6,11 group of *cdnf* wild-type brains. (**D**) Higher magnification images of TH2 10/10b (Hc) group, TH1 10 group, and TH1 5,6,11 group of *cdnf* knock-out brains. (**E**) Significant increase in TH1/TH2 immunoreactive cell number in the caudal hypothalamus area (n=8, Hc, 10/10b th population) in the *cdnf* knock-out group. (**F**) Significant decrease in TH1 immunoreactive cell number in the caudal hypothalamus area (n=8, Hc, 10 th population) in the *cdnf* knock-out group. (**G**) Significant decrease in TH1-positive cell number in the prethalamus (n=8, th1 group 5,6,11) in the *cdnf* knock-out group. (**H**) *in situ* hybridization results showing a higher intensity of *th2* signals in *th2* 10b group (Hc, caudal hypothalamus) in 8-dpf *cdnf* knock-out fish (n=8). (**I**) 5-HT immunoreactivity in 8-dpf wild-type brains (n=5). In particular, TH1+TH2 (rabbit anti-th1 and th2 antibody) recognized both zebrafish th1 and th2 dopaminergic neurons. TH1 (mouse anti-tyrosine hydroxylase antibody) specifically recognized zebrafish th1. 5-HT, rabbit anti-serotonin antibody. Data are mean ± SEM. Student’s *t*-test was used for statistical analysis. **p<0.01, ***p<0.001. Scale bar is 100μm.

### Decreased number of histaminergic neurons in *cdnf* mutant fish

We previously reported that TH2 plays a role in the regulation of histaminergic neuron development in zebrafish (29). To investigate whether the loss of functional *cdnf* causing significant TH2 upregulation affects the development of histaminergic neurons in larval *cdnf* mutants, 8-dpf dissected brains were co-stained with antibodies recognizing 1) histamine (a histamine neuron marker), and 2) HuC (a panneuronal marker) following EdU staining (a proliferation marker). A significant decrease in histaminergic neurons was found in *cdnf* mutants (Figure 5B-5C). Consistently*, in situ* hybridization of *hdc* (an enzyme converting L-histidine to histamine) showed a significant decrease in *hdc*-positive cell numbers in *cdnf* KO mutants (Figure 5D). The cell proliferation and HuC-positive pattern remained unchanged (Figure 5A, 5E, 5C, and 5G), indicating that *cdnf* deficiency directly or indirectly affects the histaminergic system, but has no effect on cell proliferation.

**Figure 5.**
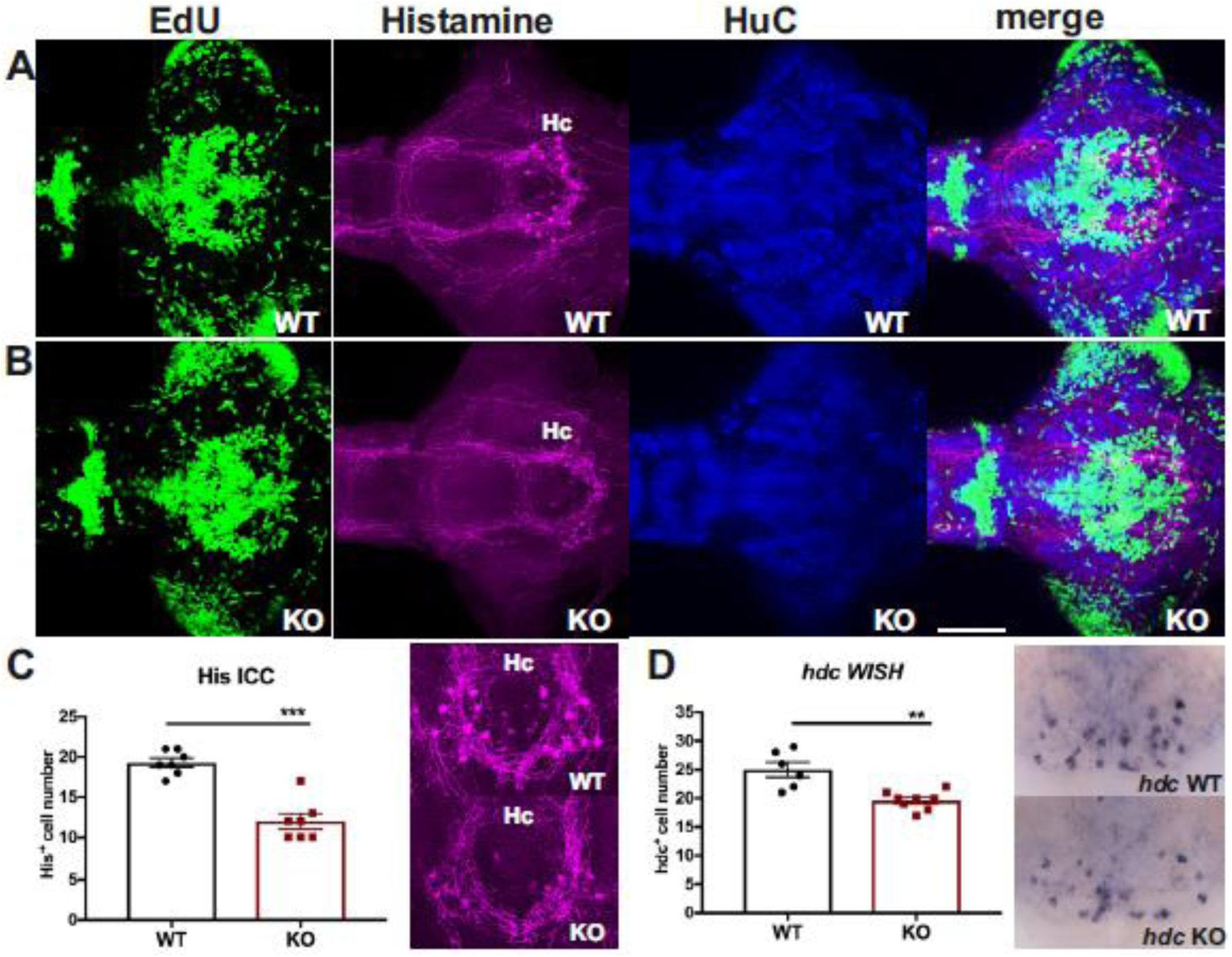
Deficient histaminergic neurons in *cdnf* mutant fish. (**A**) Co-labeling of proliferation marker (EdU), anti-histamine immunoreactivity, anti-HuC immunoreactivity, and a merged image of triple labeled stacks of an 8-dpf *cdnf* wild-type brain. (**B**) Triple labeling of an 8-dpf *cdnf* knock-out brain. (**C**) Quantification of histamine-positive cell numbers, and higher magnification images of histamine immunostaining images (n=7 *cdnf* wild-type, and n=7 *cdnf* knock-out fish). (**D**) Quantification of *histidine decarboxylase* (*hdc*) mRNA-containing cell numbers. Data are mean ± SEM of *hdc in situ* hybridization (n=8 *cdnf* wild-type, and n=8 *cdnf* knock-out fish). Student’s *t*-test was used for statistical analysis. **p<0.01, ***p<0.001. Scale bar is 100μm.

### Neurotransmitter profile by HPLC analysis

Due to the alterations of dopaminergic and histaminergic neurons in *cdnf* mutant fish, HPLC analysis was performed to measure the concentration of dopamine, norepinephrine, serotonin, histamine, and their metabolites throughout the zebrafish lifespan. No statistically significant difference was found in brain dopamine, norepinephrine, and serotonin levels, but a decrease in dopamine metabolites, DOPAC, and homovanillic acid and serotonin metabolite 5-hydroxyindoleaetic acid was found in adult *cdnf* mutant fish compared with the WT fish brains. (Table 1). A reduction in histamine level was found in 8-mpf *cdnf* mutants, consistent with a reduced expression of histaminergic marker *hdc* mRNA in 8-mpf *cdnf* KO brains (Figure 3K).

**Table 1.** HPLC analysis results of monoamine concentrations of larvae, adult brains and aging brains.

| pmole/mg protein | 19-month-old brain |  |  | 8-month-old brain |  |  | 8-dpf (8 larvae per tube) |  |  |
| --- | --- | --- | --- | --- | --- | --- | --- | --- | --- |
|  | WT (n=5) | KO (n=5) | p value | WT (n=9) | KO (n=9) | p value | WT (n=3) | KO (n=3) | p value |
| Dopamine | 22.39 ± 0.3935 | 21.14 ± 0.6298 | 0.13 | 16.52 ± 0.542 | 15.97 ± 0.433 | 0.4451 | 2.834 ± 0.1438 | 3.21 ± 0.2845 | 0.3041 |
| Norepinephrine | 58.79 ± 3.076 | 56.32 ± 2.85 | 0.5707 | 46.29 ± 1.956 | 43.3 ± 1.243 | 0.216 | 7.2 ± 0.1838 | 8.239 ± 0.56 | 0.1528 |
| DOPAC | 4.699 ± 0.3663 | 3.506 ± 0.2053 | 0.0218 | 2.764 ± 0.1298 | 2.452 ± 0.2412 | 0.2713 | 1.721 ± 0.1544 | 1.605 ± 0.0864 | 0.549 |
| Homovanillic acid | 12.48 ± 1.131 | 8.847 ± 0.1623 | 0.013 | 17.54 ± 1.768 | 8.494 ± 1.607 | 0.0016 | 2.235 ± 0.007714 | 4.195 ± 1.205 | 0.1794 |
| 3-Methoxytyramine | 1.539 ± 0.1343 | 1.343 ± 0.2165 | 0.4639 | 5.42 ± 0.6011 | 5.795 ± 0.438 | 0.6205 | 2.854 ± 0.1544 | 3.351 ± 0.4868 | 0.3855 |
| Serotonin | 23 ± 1.162 | 24.63 ± 0.6439 | 0.2543 | 16.83 ± 0.6696 | 19.43 ± 0.6398 | 0.0124 | 3.693 ± 0.157 | 3.902 ± 0.1672 | 0.4142 |
| 5-Hydroxyindoleacetic acid | 19.81 ± 0.9596 | 13.7 ± 0.4645 | 0.0004 | 15.99 ± 0.8659 | 15.08 ± 1.173 | 0.5423 | 3.769 ± 0.03767 | 5.02 ± 0.6834 | 0.1416 |
| Histamine | 9.065 ± 0.4468 | 11.33 ± 1.745 | 0.244 | 5.958 ± 0.516 | 4.285 ± 0.3379 | 0.0154 | 7.805 ± 0.854 | 5.899 ± 1.09 | 0.2406 |

### Impairments of GABAergic system

GABA acting through the GABA-A receptor in the adult brain is the major inhibitory neurotransmitter, and GABAergic neurons are widely distributed in the brain to modulate neural activity. It has been reported that MANF can potentiate presynaptic GABAergic inhibition (31). Moreover, a dual dopaminergic and GABAergic phenotype is evident in the hypothalamic areas in zebrafish (32). Dopamine signaling deficiency affects the development of GABAergic neurons in zebrafish (33). To assess whether the profound change of dopaminergic systems in *cdnf* mutant fish is associated with abnormalities in the GABAergic system, anti-GABA and anti-acetylated-alpha-tubulin antibodies (axonal alpha-tubulin marker) were used on 8-dpf dissected brains. Significantly fewer GABA-positive cells were observed in the ventral part of the posterior tuberculum (PTv) (Figure 6A-6C) and caudal hypothalamus area (Hc) (Figure 6D) of the mutant fish than in their WT siblings. The axonal tubulin pattern remained intact (Figure 6A, 6B) in *cdnf* mutants, suggesting that the loss of functional *cdnf* causes specific abnormal neurotransmitter systems rather than an overall change in neuronal organization.

**Figure 6.**
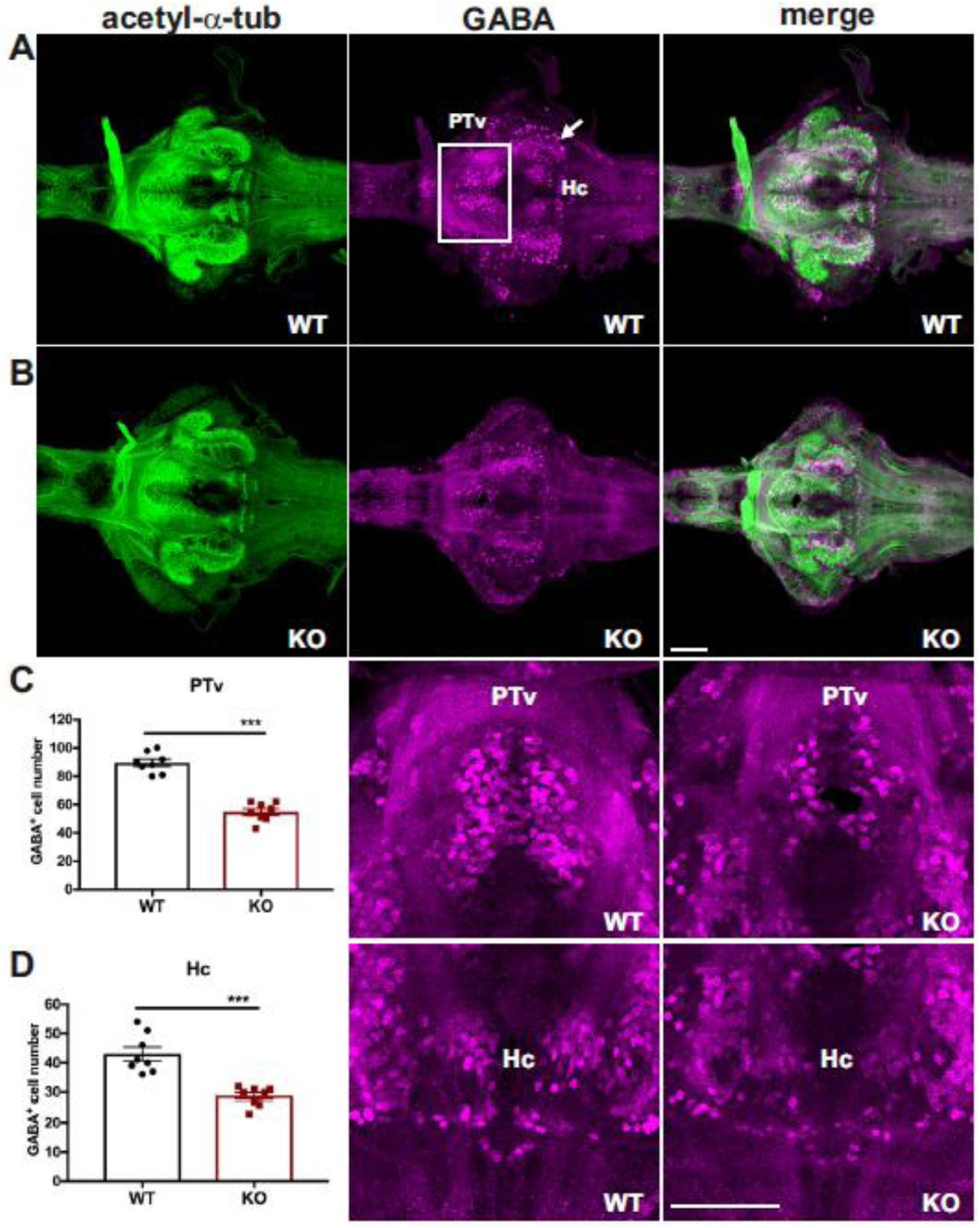
Selective impairment of GABAergic system in *cdnf* knock-out fish. (**A**) Co-labeling of anti-acetylated tubulin (showing axons) and GABA with antibody (showing GABAergic cells) of 8-dpf *cdnf* wild-type brains. (**B**) Double immunostaining of 8-dpf *cdnf* knock-out brains. (**C**) Quantification of GABA immunoreactive cell numbers in the ventral part of the posterior tuberculum (PTv), and higher magnification images of GABA-labelled cells (n=8 in each group). (**D**) Quantification of GABA immunoreactive cell numbers in the caudal hypothalamus (Hc), and higher magnification images of GABA-labelled cells (n=8 in each group). White rectangles and arrows indicate a noteworthy reduction of GABA-staining cells in the PTv and Hc area, respectively. Data are mean ± SEM. Student’s *t*-test was used for statistical analysis. ***p<0.001. Scale bar is 100μm.

GABA is converted from glutamic acid by glutamic acid decarboxylases (GADs: GAD65 and GAD67), and vesicular GABA transporter (vGAT) is responsible for uptake and storage of GABA in the vesicles in the presynaptic terminals. To study whether the decreased number of GABA-containing cells in *cdnf* mutants stems from either dysfunction of GABA synthesis or neurotransmission, we performed qPCR to determine the expression level of GABAergic markers *slc32a1/vGAT*, *gad2a/gad65* and *gad1b/gda67* in 8-dpf larvae. A significant decrease in *slc32a1/vGAT* transcripts was detected in *cdnf* mutant larvae (Figure 7A), whereas the expression level differences of *gad1b* and *gad2a* were not statistically significant (Figure 7B, 7C). Accordingly, the *in situ* hybridization results of *vGAT* in 8-dpf dissected brains showed lower expression in the ventral thalamus (VT), ventral part of posterior tuberculum (PTv) and hypothalamus (H) areas of *cdnf* KO mutants than in their WT siblings (Figure 7I). These findings suggest that the decreased number of GABA-positive cells is associated with downregulation of *vGAT* expression in the *cdnf* KO larvae.

**Figure 7.**
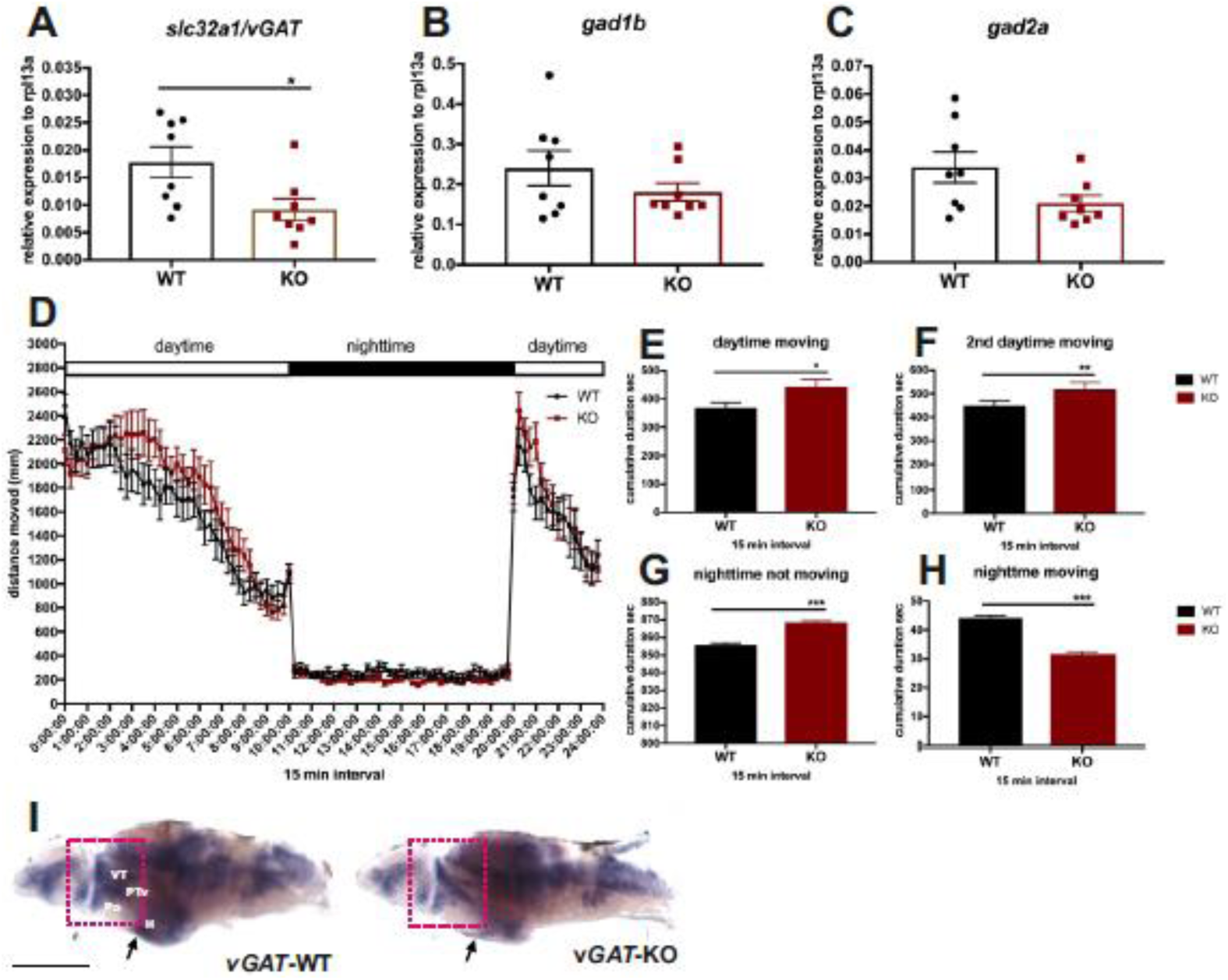
*slc32a/vGAT* downregulation in *cdnf* knock-out fish. (**A-C**) Quantification of *slc32a/vGAT*, *gad1b*, and *gad2a* mRNA expression relative to the housekeeping gene *rpl13a* in 8-dpf fish (n=8). (**D**) Locomotor activity under 14:10 (light:dark) illumination conditions. (**E**) Average cumulative duration (in s) in 15 min time bins of movement during light conditions. (**F**) Average cumulative duration of movement during the second treatment of light conditions. (**G**) Average cumulative duration of movement during dark conditions. (**H**) Duration of rest (not moving) during dark conditions, in 15 min time bins. n=16 in each genotype group. **I**, *In situ* hybridization of *vGAT* expression of 8-dpf brains (n=5). Arrows and red rectangles indicate downregulated expression of *vGAT* in the hypothalamus (H), preoptic region (Po), ventral posterior tuberculum (PTv) and ventral thalamus (VT). Data are mean ± SEM. Student’s *t*-test and one-way ANOVA analysis with multiple comparisons was used for statistical analysis *p<0.05, **p<0.01, and ***p<0.001. Scale bar is 100μm.

### Dark-flash and sleep-related behavior

Dopaminergic, histaminergic, and GABAergic circuitries play important roles in the regulation of startle response and sleep-wakefulness behavior (34). *cdnf* mutant larvae did not show impaired light/dark adaptions by the light-dark flash change (data not shown). Therefore, we examined the sleep-like locomotor behavior of 6-dpf larval fish under a 14:10 (light:dark) regime (Figure 7D, n=16 in each group). Interestingly, during the daylight period, the *cdnf* mutant larvae were more active than their WT siblings (Figure. 7E, 7F). In contrast, during the dark period, the *cdnf* mutant larvae moved significantly less than their WT siblings (Figure 7G, 7H). The abnormal locomotor activity during light and dark conditions was thus associated with the dysfunctional neurotransmission in *cdnf* mutant larvae.

### Impaired social preference in adult *cdnf* mutant fish

There is growing evidence to suggest that dysregulated neurotransmission causes neuropsychiatric disorders, some of which alter social interactions (35–37). Zebrafish are social animals and naturally tend to approach conspecifics by visual choice (38). The social preference test conducted here measures this innate tendency. To investigate the consequences of impaired dopaminergic, histaminergic, and GABAergic systems on adult fish behavior, we performed social preference analysis/visually-mediated social preference on 6-mpf fish by quantifying the amount of time each fish spent in close proximity to conspecifics (“stimulus” arena) compared to the empty “object” arena (Figure 8A). In comparison with their WT siblings, *cdnf* mutant fish spent significantly less time in the stimulus/conspecific zone (Figure 8B), but spent more time in the “distal” testing zone (Figure 8D). WT and KO fish spent similar amounts of time in the “object” zone (Figure 8C), indicating that adult *cdnf* mutant fish show less social preference for conspecifics than the WT fish.

**Figure 8.**
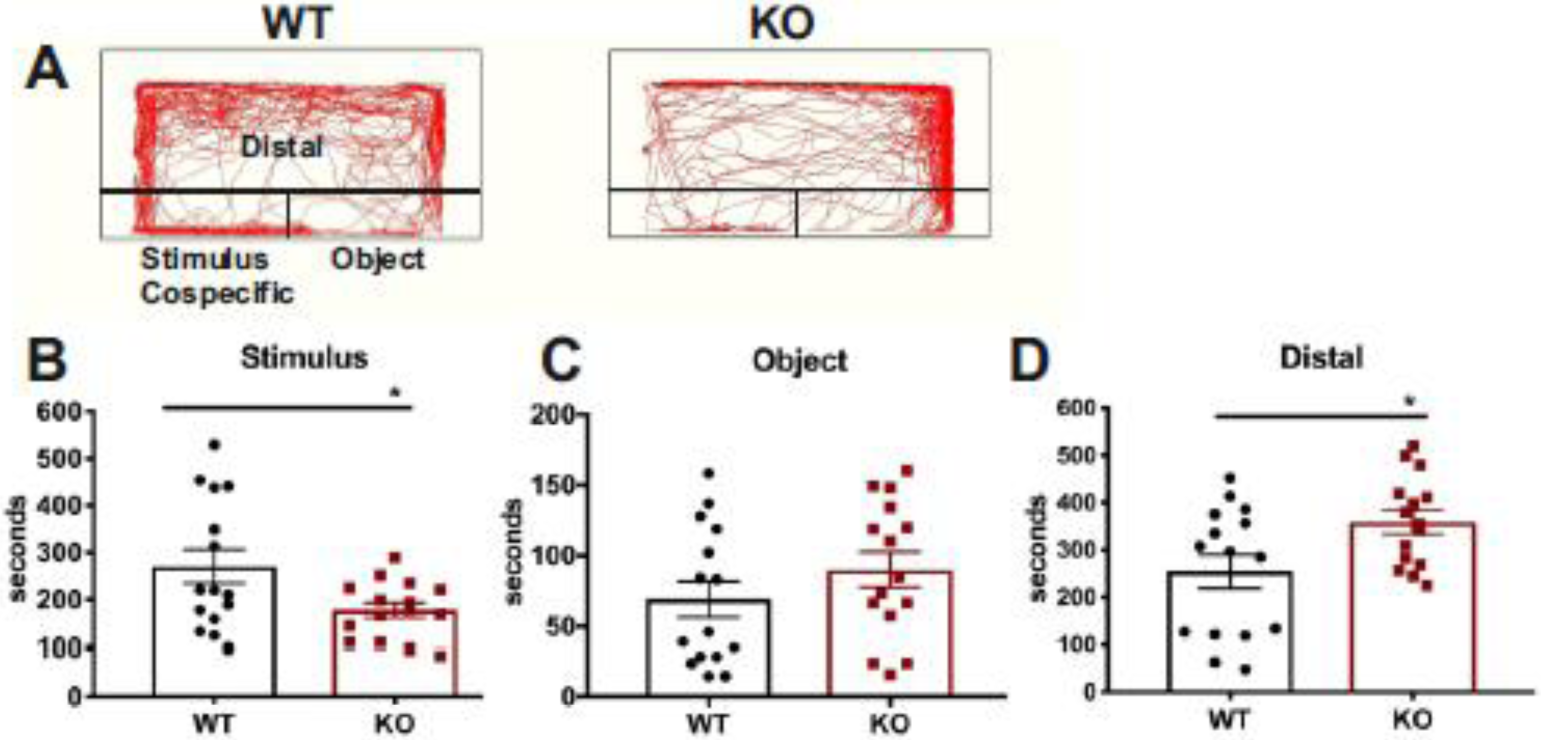
Social preference deficiency in *cdnf* knock-out adult fish. (**A**) Schemes of visually mediated social preference behavior setup and representative movement traces of *cdnf* wild-type and knock-out fish during 10 min recording intervals. (**B**) Cumulative time spent in the stimulus (conspecific) zone. (**C**) Cumulative time spent in the object zone. (**D**) Cumulative time spent in the distal zone. n=15. Data are mean ± SEM. One-way Student’s *t*-test was used for statistical analysis. *p<0.05.

### Anxiolytic behavior appeared in adult *cdnf* mutant fish

Zebrafish have a natural tendency to spend more time at the bottom of the tank when placed in a novel environment, before gradually migrating to the surface (39). We utilized a novel tank diving assay to study anxiety-related risk-taking behavior. The novel tank diving area was digitally divided into three zones. Compared with their WT siblings, *cdnf* mutants spent significantly more time exploring the top zone (Figure 9B, 9C), and less time in the bottom (Figure 9E); there was no difference in the time spent the middle zone between WT and KO siblings (Figure 9D). Representative swimming tracks are shown in Figure 9A. There was no significant difference for movement speed between WT and KO siblings, suggesting that impaired bottom-dwelling behavior was not caused by motor defects (Figure 9F).

**Figure 9.**
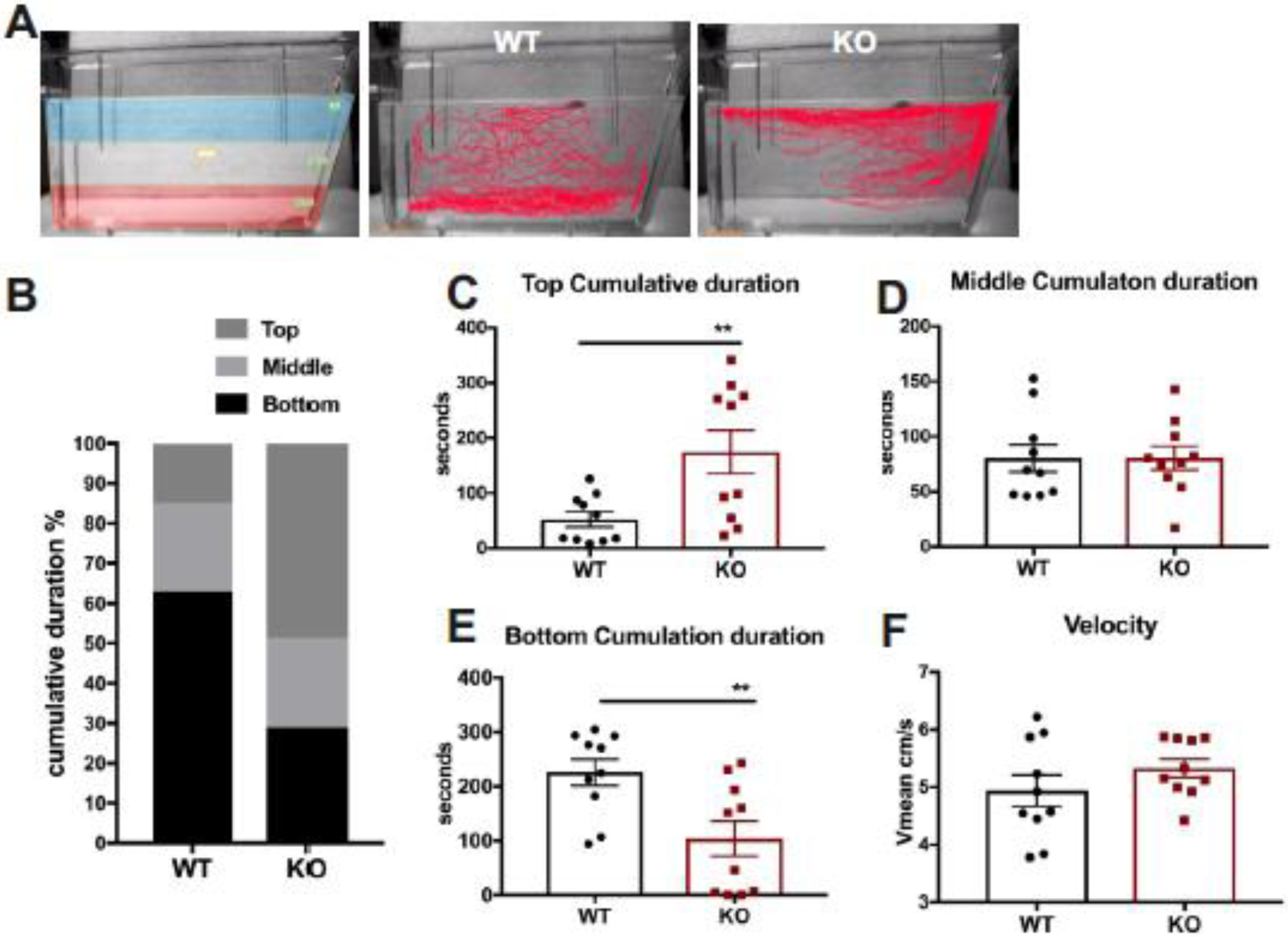
Impaired bottom-dwelling behavior in *cdnf* knock-out adult fish. (**A**) Schemes of the novel tank test with three digitized zones and representative movement traces of *cdnf* wild-type and knock-out siblings during a 6 min recording period. (**B**) Ratio of cumulative duration in the top, middle and bottom zone. (**C**) Amount of time spent in the top zone. (**D**) Amount of time spent in the middle zone. (**E**) Amount of time spent in the bottom zone. (**F**) Average velocity during the 6 min video tracking period. n=10 in each genotype group. Data are mean ± SEM. Student’s *t*-test was used for statistical analysis. **p<0.01.

### Decreased shoal cohesion in adult *cdnf* mutant fish

Zebrafish naturally swim in shoals (38). To test whether *cdnf* deficiency affects fish shoaling behavior, five 6-mpf fish (young adult group) or five 18-mpf fish (adult group) per trial were placed in a plastic cylindrical container (23 cm diameter), and monitored by video tracking for 10 min after a 15 min habituation period (n=4 trials for young adults, and n=3 trials for adults). The movement speed, average distance between the test fish and the other four shoal members, and duration of stays in proximity (the nearest inter-individual distance defined as less than 2 cm for young adults and 2.5 cm for adults) were analyzed. *cdnf* mutant fish showed higher swimming speed during 10 min locomotion activity in both age groups (Figure 10A, 10B 10E, 10F). In the young adult group, the inter-individual distance was significantly greater in the *cdnf* mutant group compared with their WT sibling group (Figure 10C). Furthermore, the time spent in proximity with shoal members was significantly shorter in *cdnf* mutant groups than in their WT siblings (Figure 10D). Similar results were obtained in the adult group. Collectively, the *cdnf* mutant fish were more hyperactive and kept at a greater distance to their neighbors, although no significant difference in time spent in proximity was observed in the adult group (Figure 10H), suggesting that lack of functional *cdnf* causes social defect phenotypes in adult zebrafish.

**Figure 10.**
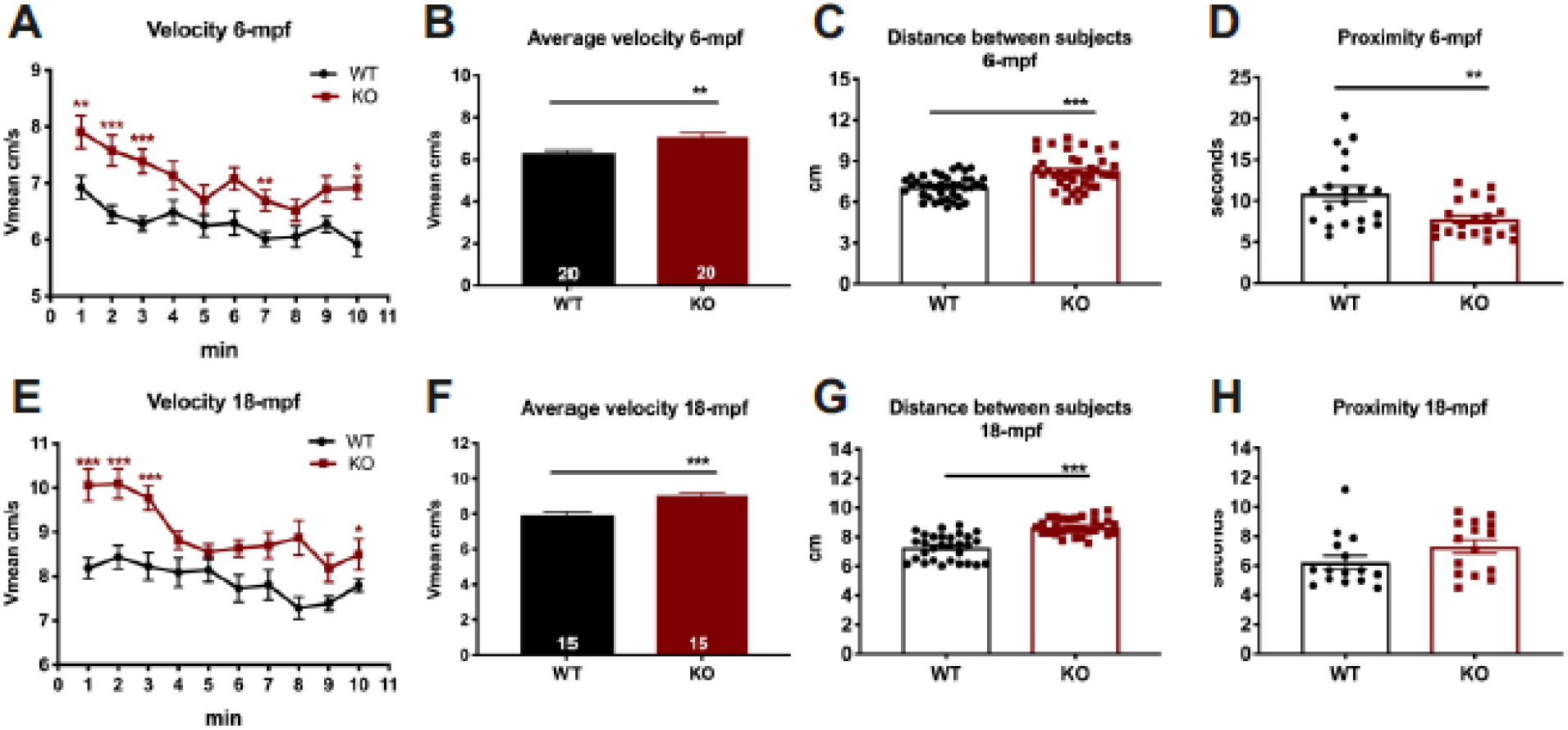
Decreased shoal cohesion in *cdnf* knock-out adult fish. The upper panel shows the shoaling behavior test on 6-mpf male *cdnf* wild-type and knock-out fish (n=5 in each trial) during a 10 min video tracking period; four trials were analyzed. (**A**) Average velocity, in 1 min time bins. (**B**) Average velocity during the 10 min video tracking period. (**C**) Average inter-individual distance of total trials. (**D**) Average time spent in the proximity (i.e. less than 2 cm) of the nearest neighbor. The lower panel shows the shoaling behavior test on 18-mpf male *cdnf* wild-type and knock-out fish (n=5 in each trial); three trials were analyzed. (**E**) Average velocity, in 1 min time bins. (**F**) Average velocity during the 10 min video tracking period. (**G**) Average inter-individual distance of total trials. (**H**) Average time spent in the proximity of closest neighbor. Data are mean ± SEM. Student’s *t*-test and one-way ANOVA analysis with multiple comparisons was used for statistical analysis. *p<0.05, **p<0.01, and ***p<0.001.

### Increased seizure susceptibility in adult *cdnf* mutant fish

We hypothesized that the impaired GABAergic phenotype found in the *cdnf* mutant fish may render the mutants more susceptible to drug-induced epileptic seizures. Pentylenetetazole (PTZ), a chemoconvulsant drug, is commonly used to induce seizures in rodents and zebrafish by inhibiting GABA-A receptor subunits (40, 41). Six-month-old fish were exposed to 10 mM PTZ (for 5 min periods, over three consecutive days) to allow analysis of the molecular consequences of PTZ-induced seizures (Figure 11A). Seizures were scored based on the description of Mussulini et al. (41). Briefly, score 3 was recorded when fish showed repetitive circular movements, score 4 included abnormal whole-body rhythmic muscular contractions, and score 5 was characterized by rigidity, loss of body posture, and sinking to the bottom of the tank. None of the tested fish died during the PTZ administration procedure. Throughout the 5 min PTZ administration, a significantly higher percentage of *cdnf* mutant fish reached score 5 compared to their WT siblings (Fig. 11B), revealing that *cdnf* mutant fish are more susceptible to PTZ-induced seizures. Moreover, the *cdnf* mutant fish showed a shorter onset latency to reach score 4 than their WT siblings, particularly on the second and third days of exposure (Figure 11C). The *cdnf* mutant fish demonstrated the longest periods of immobility (Figure 11D), but there were no significant differences between the genotypes in the total distance moved (Figure 11E).

**Figure 11.**
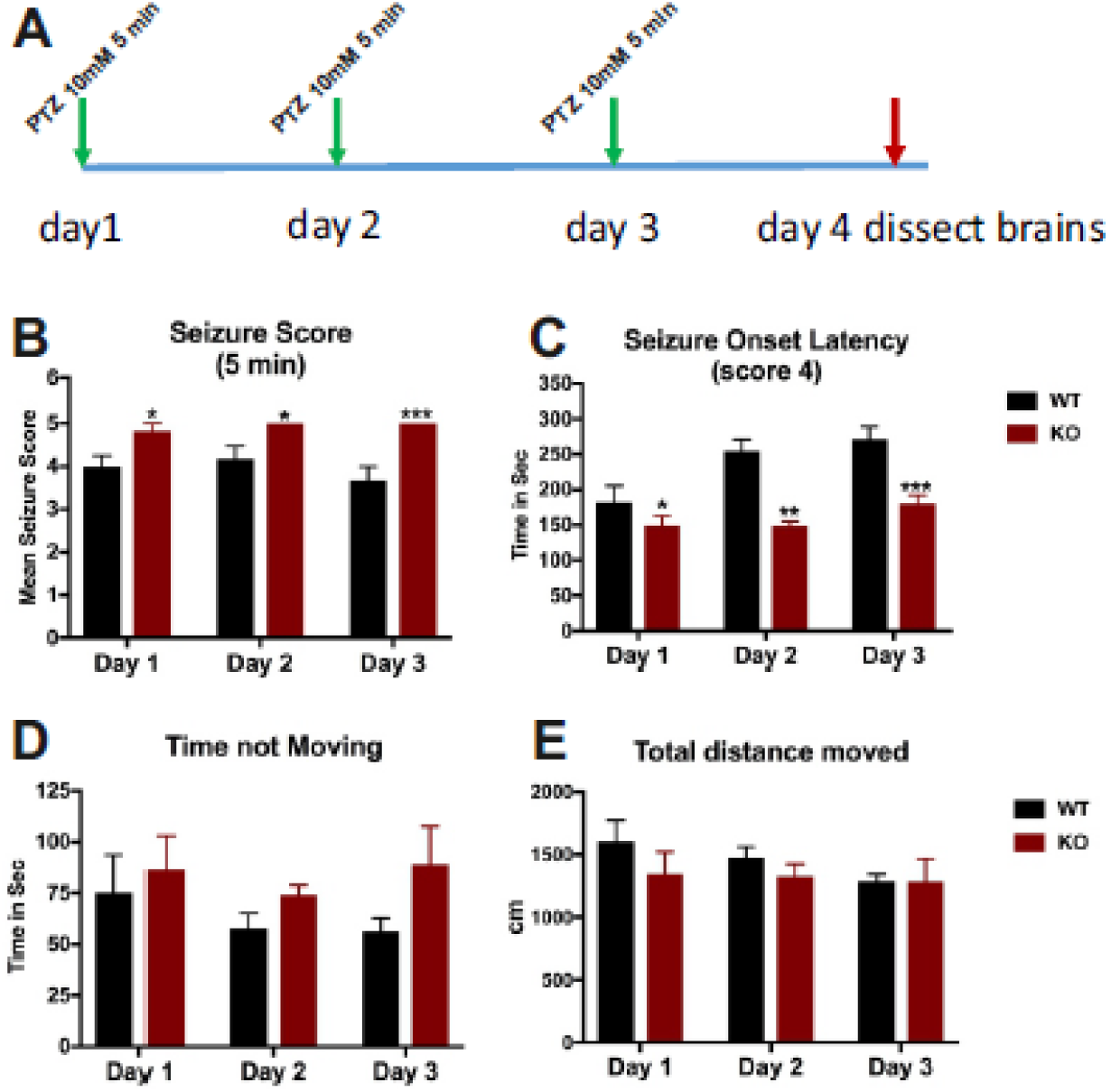
Increased seizure susceptibility in *cdnf* knock-out adult fish. (**A**) Scheme of the PTZ administration procedure. (**B**) Seizure scores after 5 min of PTZ exposure. (**C**) Seizure onset latency to score of 4. (**D**) Duration of immobility during 5 min of PTZ exposure. (**E**) Total distance travelled during 5 min of PTZ exposure. Data are mean ± SEM. Two-way ANOVA analysis with multiple comparisons was used for statistical analysis. *p<0.05, **p<0.01, and ***p<0.001.

### Gene expression in PTZ-treated *cdnf* mutant and WT fish brains

To investigate the molecular alterations in the zebrafish brain caused by PTZ treatment, qPCR was used to quantify the gene expression of *manf* (a closely related growth factor), *glial fibrillary acidic protein* (*gfap*; an astrocyte marker), and GABAergic and glutamatergic markers. We first confirmed that merely 20% of the remaining *cdnf* truncated transcript was detected in *cdnf* mutant brains, which agrees with the qPCR results (Figure 3A-3C). PTZ administration did not significantly alter the *cdnf* and *manf* mRNA expression in *cdnf* WT and mutant fish (Figure 12A, 12B). Remarkably, a dramatic downregulation of *slc17a6a/ vGlut2* (Figure 12C) and *slc32a1/vGAT* (Figure 12D) transcripts was found in *cdnf* mutants compared with WT fish. Similarly, a significantly lower expression of *gad2a/gad65* was found in *cdnf* mutants than WT fish brains (Figure 12E). *gad1b/gad67* expression was not significantly different in *cdnf* mutant and WT fish, nor was the mRNA level of those GABAergic and glutamatergic markers after PTZ treatment (Figure 12F). Expression of *gfap* was significantly lower in untreated *cdnf* mutant fish brains than in WT fish brains (Figure 12G). After PTZ treatment, a significant increase in *gfap* expression was seen in *cdnf* mutant fish brains, but not in their WT siblings (Figure 12G). The immunostaining result of zrf-1 labeled *gfap* showed that *cdnf* mutant fish revealed less extensive radial glial fibers in the hindbrain and spinal cord than their WT siblings (Figure 12H).

**Figure 12.**
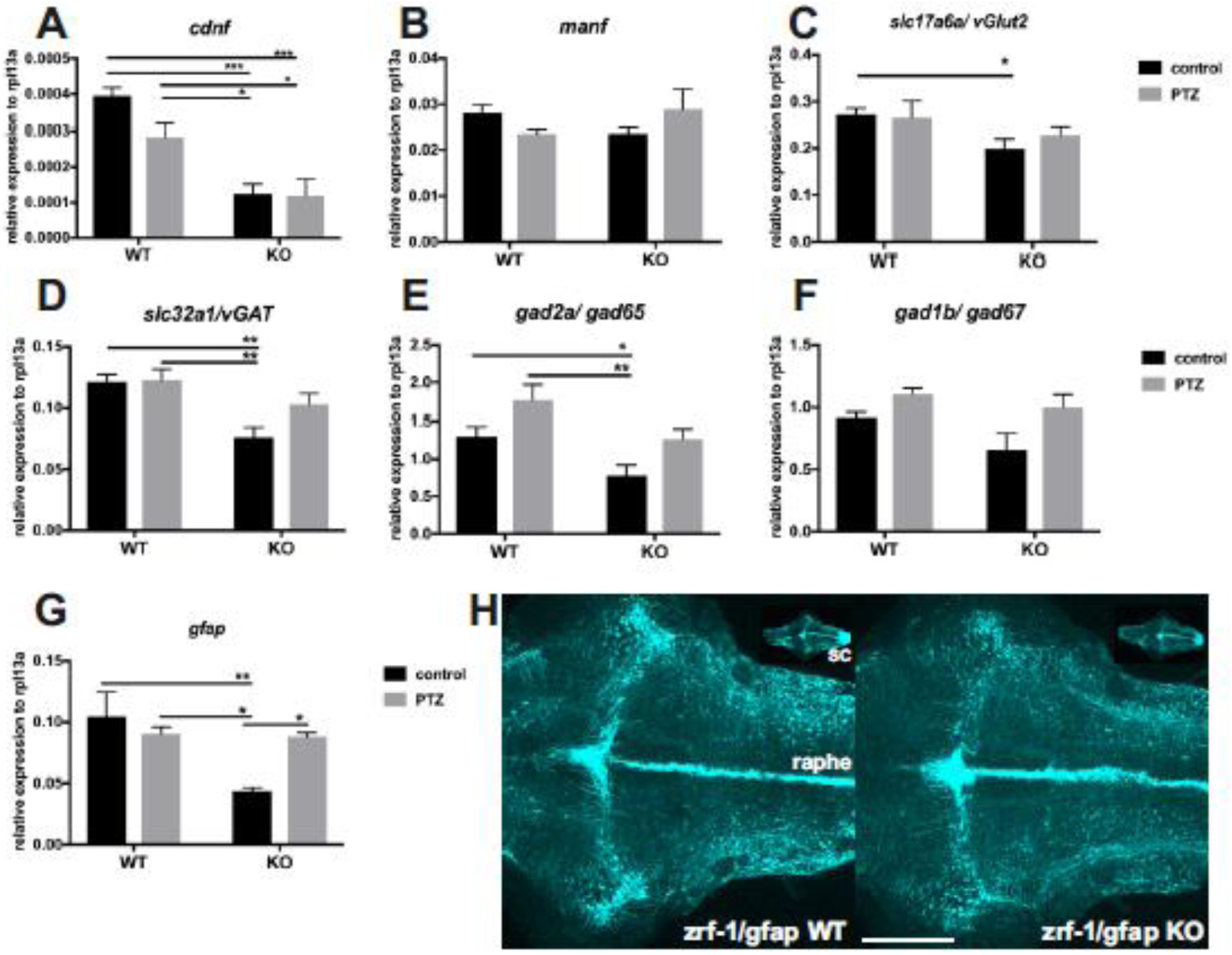
Results of qPCR analysis in PTZ-treated *cdnf* knock-out fish. Quantification of relative expression of (**A**) *cdnf*, (**B**) *manf*, (**C**) *slc17a6a/vGlut2*, (**D**) *slc32a1/vGAT*, (**E**) *gad2a/gad65*, (**F**) *gad1b/gad67*, and (**G**) *gfap* in 6-mpf brains with or without PTZ treatment. n=5 in each group. **h**, zrf-1 immunostaining images of 10-dpf *cdnf* wild-type and knock-out fish brains (n=5; sc, spinal cord). Data are mean ± SEM. Two-way ANOVA analysis with multiple comparisons was used for statistical analysis. *p<0.05, **p<0.01, and ***p<0.001. Scale bar is 100μm.

## Discussion

Despite the apparently typical general development and superficially normal behavior, zebrafish lacking CDNF displayed hyperactivity and impairments in anxiety-related behavior, social preference, and shoal cohesion. Decreased sociability and increased seizure susceptibility were associated with deficiencies in several neurotransmitter systems, including dopaminergic, GABAergic, and histaminergic neurons. Notably, there was no overall difference in whole-brain dopamine levels, but a detailed examination of the two major dopaminergic systems showed significant abnormalities in *cdnf* KO fish. These findings lend support to the hypothesis that CDNF acts as a general modulator that regulates neurogenesis and maturation of transmitter-specific neuronal types during development and throughout adulthood, rather than a regulator of only dopaminergic systems. This concept is supported by the expression pattern of *cdnf* mRNA during development and in the mature brain: it is detected in the anteroposterior axis of the ventricular zones where neurogenesis actively happens in adult zebrafish (42, 43).

There is evidence that the unfolded protein response (UPR), which is essential in the mechanisms of MANF and CDNF, is associated with the generation and maturation of CNS neurons and circuits. MANF/Armet is upregulated in various forms of ER stress (44). ATF6 is a transcription factor activated as one component of the ER stress cascade, and its conditional activation induces MANF/Armet in cardiomyocytes (45). In addition to ATF6 activation, the UPR cascade consists of activation of inositol-requiring enzyme 6 (Ire-6) and protein kinase (PKR)-like ER kinase (Perk) and is essential in nervous system development. Additionally, it supports the generation, maturation, and maintenance of CNS neurons (46, 47). For example, lack of functional BiP/GRP78 – an essential component of the UPR – disturbs development of thalamocortical connections in mice (48). Moreover, downregulation of the UPR alters generation of progenitor cells and cell fate acquisition in the developing cerebral cortex (49). In agreement with the structural similarities of MANF and CDNF (6), CDNF is also activated in the UPR. Expression of CDNF in HEK293-T cells and hippocampal neurons activates the UPR during thapsigargin-induced ER stress in both cell types, and attenuates expression of ER stress activated by the apoptotic proteins CHOP and cleaved caspase 3 (25). Interestingly, expression of CDNF that lacks the ER retention C-terminal signal QTEL abolished the protective effect in HEK293-T cells and cultured hippocampal neurons (25). This suggests that ER localization is necessary for the protective effect of CDNF. There is clear evidence that exogenously administered CDNF is also neuroprotective(11, 50). Indeed, intrastriatally infused CDNF is taken up by neurons and transported retrogradely to, for example, the cortex and substantia nigra in rats, and localized in endosomes rather than the ER (51). Although a cell surface receptor-mediated mechanism for secreted CDNF could be expected, no such mechanism has yet been identified. Moreover, all observed abnormalities are not necessarily direct consequences of a lack of CDNF, but may be secondary to the primary effects of the knockout, e.g. alterations in the GABAergic or dopaminergic systems, which are known to regulate neurogenesis and differentiation (37, 52).

Using the *cdnf* null mutant fish generated in this study, we first provide evidence that CDNF plays an important role in the regulation of developing neurotransmitter circuitries, including dopaminergic, GABAergic, and histaminergic systems. Moreover, the increased seizure susceptibility revealed by PTZ administration in adult *cdnf* KO fish may be associated with the deficiency of the GABAergic and glutamatergic systems in *cdnf* KO fish. As a consequence of this lack of functional *cdnf*, the dysregulated homeostasis of the neurotransmitter connectivity leads to the impairment of social behaviors.

### Zebrafish cdnf is structurally conserved except the ER-localizing motif at the C-terminus end

The predicted 3D structure of zebrafish cdnf was similar to the human cdnf crystal structure with dual functional domains, except for the ER-retention motif (TEEF) at the C-terminal end. This suggests that zebrafish cdnf may function similar to mammalian cdnf, but may take a non-canonical trafficking pathway shuffling between the ER and Golgi apparatus, similar to *C. elegans* MANF, although there is no evidence for this yet (53). Of the two CDNF domains, the saposin-like domain may interact with sphingolipids, like *C. elegans* MANF, acting as a cell adhesion molecule that binds to the cell membrane and activates the downstream signal cascade without binding to any receptors in different brain areas (54) to regulate the neurotransmitter systems.

### Loss of cdnf causes a dynamic alteration of dopaminergic systems in the brain

Due to the genome duplication in teleost fish (55), two TH genes (56) complimentarily expressed in the brain (28, 57, 58) are found in the zebrafish. In our *cdnf* KO mutant, the number of TH1 neurons was reduced in the prethalamus area, whereas an increased number of TH2-containing cells (teleost specific paralogous th) appeared in the hypothalamic region. The zebrafish TH1 cell population in the prethalamus is homologous to the mammalian DA population A13 in the zona incerta of the thalamus (59), which is more susceptible to neurotoxic MPTP and MPP+ injury in the zebrafish brain (60). The hypothalamic TH1 groups correspond to A12 and A14 DA groups in the arcuate and periventricular nucleus of the hypothalamus (59), respectively. In the L1CAM (neural cell adhesion molecule L1) null mice, abnormal distribution of dopaminergic neurons was evident in A12, A13 and A14 DA groups – but not the A9 group – in the substantia nigra (61). It has remained unclear whether cdnf binds to potential signaling receptors to trigger downstream signal pathways. Alternatively, it is possible that in the zebrafish brain, cdnf serves as a survival-promoting factor affecting the expression of essential regulators that could regulate the number of TH1 dopaminergic neurons but negatively control the TH2-containing cell numbers in a regional-specific fashion. Finally, we cannot exclude the possibility that loss of cdnf *per se* may induce unpredicted endoplasmic reticulum stress, oxidative stress or chronic inflammation, which could result in neurodegeneration in other more vulnerable dopaminergic populations.

### Impaired GABAergic and histaminergic system

It is evident that most of the dopaminergic neurons contain GABA as a co-transmitter in the preoptic area, prethalamus, and hypothalamus regions (32). Histamine neurons also contain GABA in all vertebrates studied thus far (62), including zebrafish (63). Moreover, we have reported that dopaminergic signaling plays a crucial role in the specification of hypothalamic neurotransmitter identity (29). The number of histaminergic neurons is determined at least in part by dopamine produced by *th2*-expressing dopaminergic neurons in zebrafish (29). The number of histaminergic neurons shows life-long plasticity in adult zebrafish through a Notch1 pathway regulated by presenilin 1 in the gamma-secretase complex (64). Dopamine activation also has a direct impact on GABAergic neuron development in zebrafish larvae (33). Consequently, a decreased number of GABA-containing cells and histaminergic neurons found in the hypothalamic area may also be caused by the dynamic alteration of dopaminergic systems, specifically the increased expression of *th2* in the hypothalamus, caused by a lack of cdnf. Nevertheless, the wide distribution of abnormal cell populations in cdnf-deficient zebrafish is more likely to derive from a direct effect on early neuronal proliferation, maturation, and transmitter specification.

### Neurotransmitter systems associated with abnormal social behaviors

The neurotransmitter phenotype in the developing and mature nervous system is regulated by genetic and environmental cues, in order to compensate for the changing homeostatic requirements and to maintain the appropriate neuronal circuits during development in nervous system function (65, 66). Proper social responses require coordinated neurotransmitter circuits in the CNS. Dysfunction of any main transmitter system is known to cause mental disorders and neurodegenerative diseases (67). The behavioral phenotype of the *cdnf* KO zebrafish is reminiscent of many neurological and psychiatric conditions, such as attention deficit disorder, autism spectrum disorder, schizophrenia, or epilepsy. The observed sensitivity to pentylenetetrazole-induced seizures may depend on an abnormal GABAergic system and low expression of vGAT, since PTZ acts through the GABA-A receptors (37, 52). The abnormal histaminergic system may be responsible for the low level of anxiety-related behavior observed in our novel tank test, since reducing histamine levels in the adult zebrafish brain by prohibiting the *hdc* inhibitor alpha-fluoromethylhistidine has the same effect (68). The decrease in *hdc* expression and number of histamine-containing posterior hypothalamic neurons found in this study may be a direct result of lack of CDNF, or a secondary effect of the increased production of dopamine by the *th2*-expressing neurons in the same cluster of cells (29).

Taken together, this study highlights the novel and broad role of CDNF in shaping the neurotransmitter circuits in the zebrafish brain. Although the *cdnf* KO fish are superficially normal, the altered transmitter networks produce a range of abnormal behaviors that resemble some human neuropsychiatric conditions, including schizophrenia. Indeed, one study has already shown an association between one SNP/haplotype in the human cdnf gene and schizophrenia characterized by negative symptoms in the Han Chinese population (69). Interestingly, ER quality control of protein processing is known to be associated with schizophrenia (70).

## Methods

### Zebrafish Maintenance

Zebrafish were obtained from our wild-type (Turku) line that has been maintained in the laboratory for more than a decade (29, 71, 72). Fish were raised on 14:10 (light:dark) cycles at 28°C and staged in hours post-fertilization (hpf), days post-fertilization (dpf), or months post-fertilization (mpf) as previously described (73). The permits for all experiments were obtained from the Office of the Regional Government of Southern Finland, in agreement with the ethical guidelines of the European convention.

### RNA isolation and cDNA synthesis

Total RNA was extracted from ten pooled larval fish or one dissected adult brain per sample (RNeasy mini Kit; Qiagen, Valencia, CA, USA) for quantitative RT-PCR (qRT-PCR). To synthesize cDNA, 2μg of total RNA was reverse-transcribed using SuperScriptTM III reverse transcriptase (Invitrogen, Carlsbad, CA, USA) with random hexamer primers (Roche Diagnostics, Germany) according to the manufacturer’s instructions.

### *In Situ* Hybridization

Whole-mount *in situ* hybridization (WISH) was performed on 4% paraformaldehyde (PFA)-fixed embryos and dissected brains as described earlier (28). Antisense and sense digoxigenin (DIG)-labelled RNA probes were generated using the DIG RNA labelling kit (Roche Diagnostics, Germany) following the manufacturer’s instructions. The WISH procedure was followed according to the protocol described by Thisse & Thisse (74), with slight modifications. Briefly, the prehybridization and hybridization steps were conducted at 60°C for all riboprobes. The specificity of the anti-sense ribosprobe *hdc* and *th2* have been described earlier (28, 29). The cloning primers for the open-reading frames of *cdnf* and *vGAT* cDNA are listed in Table 1. *In situ* hybridization signals were detected with sheep anti-digoxigenin-AP Fab fragments (1:10,000; Roche Diagnostics, Germany). The color staining was carried out with chromogen substrates (nitro blue tetrazolium and 5-bromo-4-chloro-3-indolyl-phosphate) and incubated in the dark at room temperature. Stained samples were embedded in 80% glycerol and visualized under brightfield optics using a Leica DM IRB inverted microscope with a DFC 480 charge-coupled device camera. Z-stacks were processed with Leica Application Suit software, with the multifocal algorithm to identify the gene expression patterns (16).

### Quantitative real-time PCR (qPCR)

qPCR was performed with a LightCycler^®^ 480 instrument (Roche, Mannheim, Germany) using the Lightcycler^®^480 SYBR Green I master mix (Roche, Mannheim, Germany) according to the manufacturer’s instructions. Primers for amplification were designed by Primer-BLAST (NCBI) and are listed in Table 2. Two housekeeping genes, β-*actin* and *ribosomal protein L13a* (*rpl13a*) were used as reference controls. All primer sets were confirmed to amplify only a single product of the correct size. Cycling parameters were as follows: 95°C for 5 min, followed by 45 cycles of 95°C for 10 s, 60°C for 15 s, and 72°C for 20 s. Fluorescence changes were monitored with SYBR Green after every cycle. Dissociation curve analysis was performed at the end of the cycles (0.1°C per s increase from 60°C to 95°C with continuous fluorescence readings) to ensure that only a single amplicon (single melting peak) was obtained. All reactions were performed in duplicates, and at least three individual biological replicates were used (sample numbers indicated in figure legends). Duplicate quantification values were analyzed with the LightCycler 480 software. The data were calculated by the comparative method, using Ct values of β-*actin* and *rpl13a* as a reference control (75). Since the changes of relative gene expression showed the same trend when normalized to the different housekeeping genes (data not shown), only the results from *rpl13a* are presented.

**Table 2.**
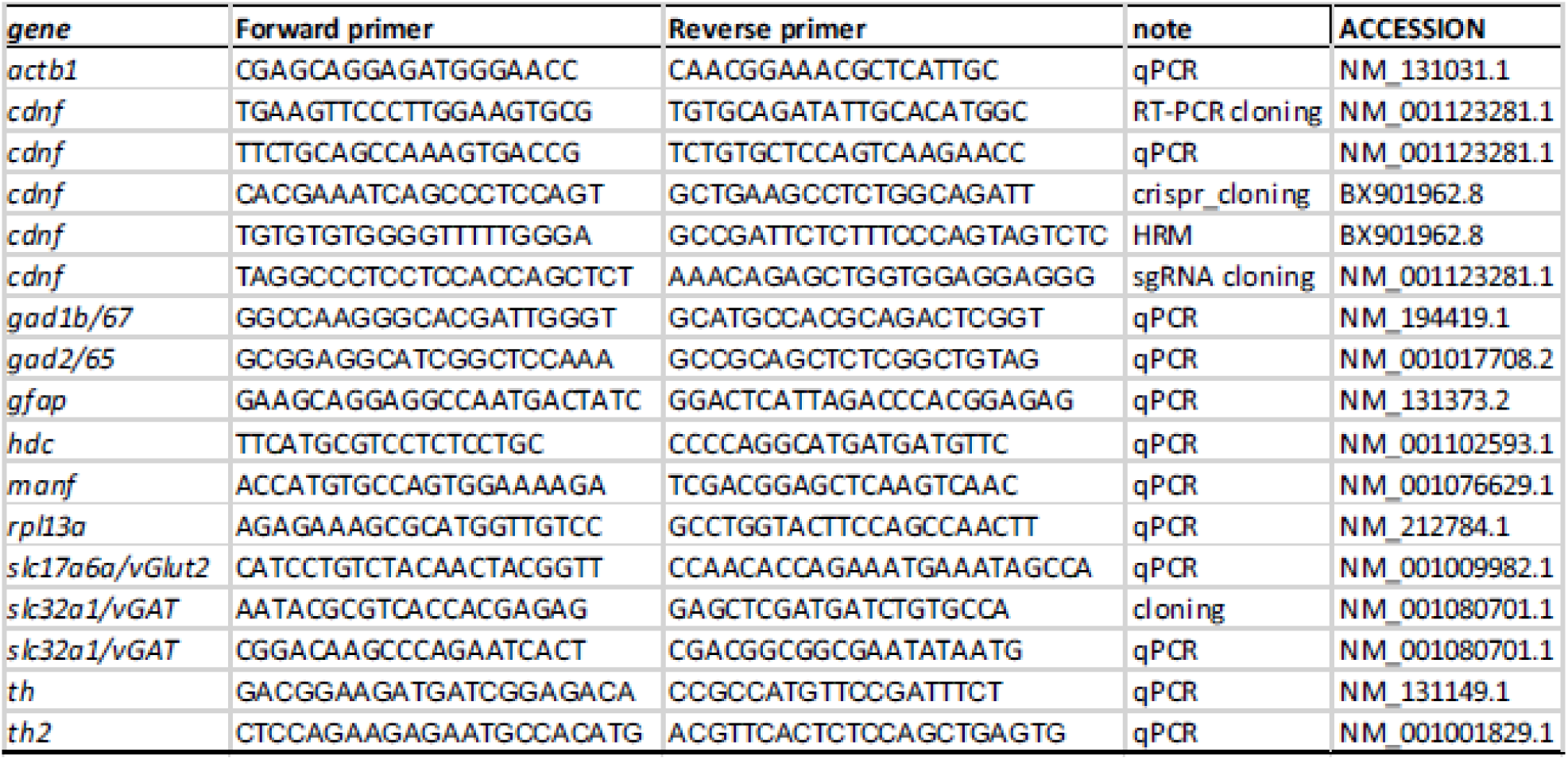
List of primers used in this study.

### Establishing zebrafish mutants

CRISPR/Cas9-genome edited fish were generated in our Turku wild-type strain, based on the description of Hwang et al. (76). To avoid off-target genomic mutagenesis effects, targeting sites were selected with a minimum of three mismatches in the genome as predicted by the CHOPCHOP software (http://chopchop.cbu.uib.no). The sequence-specific sgRNA template was generated in a pDR274 vector (Addgene Plasmid #42250; oligo sequence listed in Table 2). The sequences of the modified plasmids were verified by Sanger sequencing. sgRNAs were transcribed from linearized template plasmids (Ambion MEGAscritp), and Cas9 mRNA was transcribed *in vitro* from linearized template plasmid pMLM 3613 (Addgene Plasmid #42251). A mixture containing approximately 300ng/μl Cas9 mRNA and 20ng/μl sgRNA was injected into fertilized eggs at the one-cell stage. To verify the mutation efficiency of Cas9-sgRNA genome editing, the injected eggs were collected at 24 hpf. DNA was extracted from individual embryos and non-injected controls. PCR amplicons encompassing the targeted sites were amplified and analyzed via Sanger sequencing and high-resolution melting (HRM) analysis. Mutations were recognized as multiple sequencing peaks at the sgRNA target site. When the mutation efficiency was over 50%, the remaining Cas9/sgRNA-injected embryos were raised to adulthood and out-crossed to the wild-type fish to generate F1 progeny. To collect the tail biopsies, 3-dpf embryos were anesthetized with 0.02% Tricaine. The tip of the caudal fin within the pigment gap was removed using a microscalpel, and each larva was placed in an individual well of a 24-well plate with fresh embryonic medium until 5 dpf. We have consistently achieved 100% survival rate from the beginning of this procedure to adulthood. F1 genotyping was done by HRM assays and DNA sequencing. To identify mutated alleles from single embryos, each target locus was PCR amplified from individual genomic DNA with gene-specific primers (Table 2). PCR products were then cloned and sequenced. Mutated alleles were identified by comparison with the wild-type sequence. Heterozygous (HET) F1 siblings carrying the same mutations were pooled in one tank and raised to adulthood. Due to the sex imbalance in the F1 generation of *cdnf* HET fish, F1 male HET fish were outcrossed to Turku female wild-type fish; female mutants were obtained from the resulting F2 progeny. Genotyping of the F2s was done as described for the F1 generation.

### Fin clipping and genomic DNA extraction of adult zebrafish and 3 dpf larvae

To lyse genomic DNA, individual tail clippings were incubated in 50μl lysis buffer (10mM Tris-HCl pH8.3, 50mM KCl, 0.3% Tween-20 and 0.3% NP-40) at 98°C for 10 min, followed by incubation on ice for 2 min. 1μl of Proteinase K (20mg/ml) was added to remove protein, and the mixture was incubated at 55°C for at least 4 h. To inactivate Proteinase K activity, samples were incubated at 98°C for 10 min and quenched on ice. To detect indel mutations, HRM curve acquisition and analysis was performed. Primers flanking the mutation site were designed using Primer-BLAST (https://www.ncbi.nlm.nih.gov/tools/primer-blast; sequences are listed in Table 2). The HRM analysis was performed on a LightCycler® 480 instrument (Roche) using the following reaction mixtures: 1× LightCycler 480 HRM master mix (Roche), 2mM MgCl_2_, and 0.15μM primer mixtures. The PCR cycling protocol was as follows: one cycle of 95°C for 10 min; 45 cycles of 95°C for 10 s, 60°C for 15 s, 72°C for 20 s, and melting curve acquisition; one cycle of 95°C for 60 s, and 40°C for 60 s. PCR products were denatured at 95°C for 60 s, renatured at 40°C for 60 s, and melted at 60°C to 95°C with 25 signal acquisitions per degree. Melting curves were generated over a 65– 95°C range. Curves were analyzed using the LightCycler® 480 gene-scanning software (version 1.5) according to the manufacturer’s instructions (Roche Diagnostics Ltd., Switzerland). To identify deviations of the curves indicative of sequence mutations, a three-step analysis was performed using the Gene Scanning program (Roche) as follows: (1) Normalizing the raw melting-curve data by setting the initial fluorescence uniformly to a relative value of 100% and the final fluorescence to a relative value of 0%. (2) Determining the temperature threshold at which the entire double-stranded DNA was completely denatured. (3) Further analyzing the differences in melting-curve shapes (threshold setup 0) in order to cluster the melting curves with similar shapes into the same groups. Those with analogous melting curves were characterized as the same genotype.

### Analysis of catecholamines and histamine by high performance liquid chromatography (HPLC)

For each sample, ten 8-dpf larvae were pooled into a group. The dissected brains of 8-mpf or 18-males were flash-frozen in liquid nitrogen and individually homogenized with sonication in 150μl of 2% perchloric acid. After centrifugation, 10μl of supernatant was assessed for monoamine concentration measurement. The detection details are described in Sallinen et al. (60, 77). The results were normalized to the total protein concentration of each sample, which was measured using the Pierce^©^ BCA Protein Assay Kit. The HPLC analysis was carried out as a blinded experiment.

### Immunocytochemistry

Immunostaining was performed on zebrafish fixed in 2% PFA or 4% 1-ethyl-3 (3-dimethylaminopropyl)-carbodiimide (EDAC, Carbosynth, Berkshire, UK). For larvae older than 5 dpf, fixed brains were dissected to enhance antigen presentation and improve image quality. Antibody incubations were carried out with 4% normal goat serum and 1% dimethyl sulfoxide (DMSO) in 0.3% Triton X-100/ phosphate buffered saline (PBS) for 16 h at 4°C with gentle agitation. Primary antibodies were rabbit anti-histamine 19C (1:5,000; (72, 78)), rabbit anti-TH2 antibody (1:2000; (30)), rabbit anti-serotonin antibody (1:1000; S5545, Sigma, St. Louis, MO, USA), and anti-tyrosine hydroxylase (TH1) monoclonal mouse antibody (1:1000; Product No 22941, Immunostar, Husdon, WI, USA). The specificities of the anti-histamine, commercial anti-mouse monoclonal TH, anti-rabbit-TH2 and anti-serotonin antibodies have been verified previously (71). The following secondary antibodies were applied: Alexa Fluor® 488 and 568 anti-mouse or anti-rabbit IgG (1:1000; Invitrogen, Eugene, OR, USA).

### Immunocytochemistry following EdU proliferation labeling

To detect the proliferating S-phase dividing cells, the Click-iT^TM^EdU Alexa Fluor 488 imaging kit (Molecular Probes) was used following the manufacturer’s instructions, with minor modifications. Briefly, 5-dpf larvae were incubated in 0.5mM EdU/E3 buffer (zebrafish embryonic medium; 5 mM NaCl, 0.44 mM CaCl_2_, 0.33 mM MgSO_4_, and 0.17 mM KCl) with 1% DMSO for 24 h at 28°C. Labelled samples were transferred back to E3 medium for 30 min and fixed in 4% EDAC/PB buffer pH 7.0 overnight at 4°C with gentle agitation. The skin and lower jaw of the fixed specimens were removed in order to enhance sample penetration. Dissected brains were incubated with rabbit anti-Histamine 19C antibody (1:5000) and mouse anti-HuC antibody (1:1000). The secondary antibody Alexa Fluor®568 anti-rabbit IgG and Alexa Fluor®633 anti-mouse IgG (1:1000; Invitrogen, Eugene, OR, USA) were applied. After immunostaining, labelled specimens were fixed in 4% PFA/PB for 20 min at room temperature, and then incubated in 1× Click-iT EdU cocktail with the green-fluorescent Alexa Fluor® 488 azide dye for one hour in the dark at room temperature. After removing the reaction cocktail and rinsing in 1×PBST (phosphate-buffered saline and 0.25% Triton X-100) three times for 10 min, samples were mounted in 80% glycerol/PBS for confocal microscopy imaging.

### Imaging

Brightfield images were taken with a Leica DM IRB inverted microscope with a DFC 480 charge-coupled device camera. Z-stacks were processed with Leica Application Suite software and Corel DRAW 2018 software (28). Immunofluorescence samples were examined using a Leica TCS SP2 AOBS confocal microscope. For excitation, an Argon laser (488 nm), green diode laser (561 nm), and red HeNe laser (633 nm) were used. Emission was detected at 500–550 nm, 560–620 nm, and 630–680 nm, respectively. Cross-talk between the channels and background noise were eliminated with sequential scanning and frame averaging as previously described (60). Stacks of images taken at 0.2– 1.0 μm intervals were compiled, and the maximum intensity projection algorithm was used to produce final images with Leica Confocal software and Imaris imaging software (version 6.0; Bitplane AG, Zurich, Switzerland). Cell numbers were counted in each 1.0 μm optical slice using ImageJ 1.52b software (National Institutes of Health, Bethesda, USA). All cell counts were performed by an investigator blinded to the sample type.

### Dark-light flash and sleep behavior test for larval zebrafish

Behavioral trials were done between 11:00 and 16:00. For larval locomotion tracking, 6-dpf zebrafish larvae were individually placed in a 24-well culture dish well containing approximately 1.5mL of E3 medium. The light level was set to approximately 330 lux based on the setting of Puttonen et al. (79). Before each trial, the larvae were habituated in the observation chamber for 10 min, followed by a 30 min locomotion tracking period with the lights on. A dark-light flash response was induced by switching off the lights for 2 min, then turning them back on for 2 min. One experiment consisted of three subsequent periods of white lights on and white lights off. Locomotor activity was monitored for one day with continuous illumination by infrared lights while white light remained on from 12:00 to 22:00 on the first day and from 8:00 to 12:00 on the next day. Locomotion response was monitored at room temperature using the Daniovision system (Noldus, Wageningen, The Netherlands). Video tracking was analyzed by EthoVision XT 8.5 locomotion tracking software (Noldus, Wageningen, The Netherlands).

### Social interaction assay

The visually mediated social preference test was based on the setup of Baronio et al. (80). Briefly, an acrylic apparatus (29 cm length × 19 cm height × 29 cm width) was divided into three arenas by two acrylic partitions. A rectangular compartment in the middle was the testing arena, referred to as the “distal” zone; to one side, the conspecific compartment housed a group of six fish, referred to as the “stimulus” zone; the other adjacent compartment was filled with stones and plant mockups, referred to as the “object” zone. A single 6-mpf adult was placed in the testing arena to allow exploration and analysis of place preference. All experimental fish were raised in a social environment. All behavioral tests were performed between 11:00 and 16:00, and video-recorded from above the tank for 6 min. To quantify social preference, the videos were analyzed with the EthoVision XT 8.5 locomotion tracking software (Noldus, Wageningen, The Netherlands), and the amount of time each test fish spent in the proximity of each compartment was quantified.

### Novel diving tank assay

The novel tank assay was performed based on Cachat et al. (81). One day before the experiment, 6-mpf male adult fish with home tanks (19 cm x 34 cm x 21 cm) were habituated in the behavior testing room. In each trial, one fish was placed in a transparent tank (24 cm × 14.5 cm × 5 cm) with 1 L of fish system water. All behavioral tests were performed between 11:00 and 16:00, and video-recorded from the side of the tank for 6 min, using a Basler acA1300-60gm industrial CCD video camera. We performed a three-compartment novel tank test, with digitized divisions between top, bottom, and middle virtual zones. The time spent in each zone was quantified using EthoVision XT 8.5 software. Fish were returned to their home tanks immediately after the test.

### Shoaling assay

Five 6-mpf or five 18-mpf male fish per cohort were placed in a round white polyethlene plastic flat-bottomed container (23 cm height, 23 cm diameter) with 2 L of fish system water (5.0 cm depth) based on the description of Green et al. (82). Prior to testing, fish were habituated for 15 min followed by video recording for 10 min with a camera at a fixed height (60 cm) from the top of the container. All videos were analyzed with EthoVision XT 8.5 software, using the default setting (the center-point detection of the unmarked animals). The mean of the inter-fish distance (defined as distance between the body center of every member of the shoal) (82) was quantified from the average data from all trials (n=4 trials for the 6-mpf fish, and n=3 trials for the 18-mpf fish). The proximity duration (in s) was defined as the average duration of time a fish stayed close to the shoal fish (i.e. within 2 cm for the 6-mpf fish or 2.5 cm for the 18-mpf fish). The misdetection rate of the video-tracking software was less than 1%. All behavioral trials were done between 11:00 and 16:00.

### Seizures induced by pentylenetetrazole (PTZ) in adult zebrafish

6-mpf fish were individually exposed to 10 mM PTZ in a 1 L tank (24 cm length × 5 cm width × 14.5 cm height) for 5 min on three consecutive days, in order to induce experimental seizures based on the description of Duy et al. (83). Epileptic seizure stage scores were assigned according to Mussulini et al. (41). After PTZ administration and seizure analysis, treated fish were transferred to a clean tank for one day until they recovered (i.e. seizure score = 0). They were then sacrificed by decapitation after cold-shock, and brains were dissected for RNA extraction. The PTZ concentration and the exposure period were selected and optimized based on our pilot study, which aimed to determine the shortest time of PTZ exposure that induces a seizure of score V (including fish falling to the bottom of the tank and loss of the body posture for 1–2 s), but allowing full recovery after three daily exposures(83). Control fish were subjected to the same procedure, but exposed to only clean system water.

### Statistical analysis

Data analysis was performed by GraphPad Prism software (version 7; San Diego, CA, USA). p-values were generated by one-way analysis of variance (ANOVA) for multiple comparisons using Tukey’s multiple comparison test, two-way ANOVA for multiple comparisons, and Student’s unpaired *t*-test for comparison of two groups. Data were presented as mean ± SEM. Statistical significance was considered at p-value <0.05.

### Animal experiment permits

Permits for this study were granted by the Regional State Administrative Agencies (ESAVI/6100/04.10.07/2015 and ESAVI/13090/2018).

## Author contributions

YCC designed the study, conducted and performed experiments, interpreted data and wrote the manuscript. DB and SS performed experiments, acquired data and assisted preparation of the manuscript. SA provided vGAT materials. PP conceived and designed the study and wrote the manuscript.

## Acknowledgements

This study was supported by grants from the Jane and Aatos Erkko Foundation, Sigrid Juselius Foundation, Magnus Ehrnrooth’s Foundation and Finska Läkaresällskapet. We thank Mr. Henri Koivula (BSc), Ms Riikka Pesonen (BSc), and Ms. Noora Hellen (MSc) for expert technical help. We thank Dr. Mart Saarma, Dr. Esa Korpi and Dr. Marnie Halpern for constructive comments on the manuscript.

## Notes

The authors have declared that no conflict of interest exists

#### Summary of Updates

The abstract is shortened to emphasize the main points and clinical relevance of the study, because CDNF is now in clinical trials. Figure legends are now on the same page as the figures for clarity.

